# Schengen-pathway controls spatially separated and chemically distinct lignin deposition in the endodermis

**DOI:** 10.1101/2020.10.07.329664

**Authors:** Guilhem Reyt, Priya Ramakrishna, Isai Salas-González, Satoshi Fujita, Ashley Love, David Tiemessen, Catherine Lapierre, Kris Morreel, Monica Calvo Polanco, Paulina Flis, Niko Geldner, Yann Boursiac, Wout Boerjan, Michael W. George, Gabriel Castrillo, David E. Salt

## Abstract

Lignin is a complex polymer precisely deposited in the cell wall of specialised plant cells, where it provides essential cellular functions. Plants coordinate timing, location, abundance and composition of lignin deposition in response to endogenous and exogenous cues. In roots, a fine band of lignin, the Casparian strip encircles endodermal cells. This forms an extracellular barrier to solutes and water and plays a critical role in maintaining nutrient homeostasis. A signalling pathway senses the integrity of this diffusion barrier and can induce over-lignification to compensate for barrier defects. Here, we report that activation of this endodermal sensing mechanism triggers a transcriptional reprogramming strongly inducing the phenylpropanoid pathway and immune signaling. This leads to deposition of compensatory lignin that is chemically distinct from Casparian strip lignin. We also report that a complete loss of endodermal lignification drastically impacts mineral nutrients homeostasis and plant growth.

## INTRODUCTION

Lignin is a phenolic polymer and is one of the main components of secondary-thickened cell wall in vascular plants. Its chemical properties give strength, stiffness and hydrophobicity to the cell wall. Lignin provides mechanical support, modulates the transport of water and solutes through the vascular systems, and provides protection against pathogens (1, 2). Lignin polymerisation occurs through oxidative coupling of monolignols and other aromatic monomers (3, 4). The monolignols, that is *p*-coumaryl, coniferyl, and sinapyl alcohols are synthesized from the amino acid phenylalanine through the phenylpropanoid pathway. They are then polymerised into lignin to form the *p*-hydroxyphenyl (H), guaiacyl (G), and syringyl (S) subunits of the lignin polymer. Lignin composition and abundance are highly variable among and within plants species, tissues, cell types and can be modulated by environmental cues (1).

In roots, large amounts of lignin is deposited in the xylem vessels, an important component of the vascular system (5, 6). Yet, lignin is also deposited in the endodermal cells surrounding the vascular tissues, for Casparian strip (CS) formation (7). Both the vascular system and the CS play a critical role for water and mineral nutrient uptake from the soil and their transport toward the shoot (8-10). In *Arabidopsis thaliana*, the composition of lignin monomers in CS and xylem is similar with a strong predominance of G-unit (>90%) (7). However, the machinery required for CS lignification appears to be distinct from that needed for xylem lignification (6, 11).

The deposition of the CS in the endodermal cell wall prevents the apoplastic diffusion of solutes between the outer and inner tissues of the root, forcing solutes to pass through the symplast of endodermal cells (8). CS lignin encircles each endodermal cell, forming a bridge between them. This precise lignin deposition is defined by the presence of the transmembrane Casparian strip domain proteins (CASPs) (12), peroxidases (13, 14) and the dirigent-like protein ESB1 (15). The expression of this lignin polymerisation machinery is tightly controlled by the transcription factor MYB36 (16, 17). A surveillance mechanism for CS integrity, called the Schengen-pathway, boosts CS deposition and is necessary for CS fusion and sealing of the extracellular space (apoplast) (18). This pathway involves vasculature-derived peptides CASPARIAN STRIP INTEGRITY FACTORS 1 and 2 (CIF1 and 2) (19, 20) and their perception by the leucine-rich repeat receptor-like kinase (LRR-RLK) called SCHENGEN3 (SGN3, also called GSO1). Their interaction triggers a cascade of signalling events mediated by kinases, that involves SGN1, and the activation of the NADPH oxidase RBOHF (SGN4) leading to ROS production, necessary for lignin polymerisation (14, 18, 21). These kinase signalling events occur on the cortex-facing side of the CS and mediates the transition from a discontinuous CS with islands of lignin into a continuous CS with its characteristic ring of lignin that seals the apoplast (18). Once the CS is sealed, CIF peptide diffusion is blocked and the Schengen-pathway becomes inactive. In mutants with an impaired CS, such as *esb1* and *myb36*, the Schengen-pathway is constitutively activated due to a constant leak of the CIF(s) peptides through the CS region (12, 15, 17, 18, 22). This induces a compensatory lignification in the cell corners and suberisation of endodermal cells. However, the role of this compensatory lignin and the mechanism controlling its deposition are not fully understood.

Here, we demonstrate that the constitutive activation of the Schengen-pathway induces the deposition of a compensatory lignin in the corners of endodermal cells chemically distinct from CS lignin. We characterised this lignin and found commonalities with stress- and pathogen-response lignin, which has a high content of H subunit. Furthermore, we demonstrate that this cell corner lignification is preceded by a transcriptional reprogramming of endodermal cells, causing a strong induction of the phenylpropanoid pathway and a significant inactivation of aquaporin expression. Our findings also establish that the activation of the Schengen-pathway to compensate for a defective CS is of critical importance for plants to maintain their mineral nutrients homeostasis and water balance.

## RESULTS AND DISCUSSION

### SGN3 and MYB36 control two pathways leading to different endodermal lignification

In order to disentangle the role of MYB36 and SGN3 in controlling endodermal lignification, we generated the double mutant *sgn3-3 myb36-2*. We analysed the endodermal accumulation of lignin in the double mutant *sgn3-3 myb36-2*, and the corresponding single mutants *sgn3-3* and *myb36-2* (Fig. 1A-C). In the early stage of endodermal differentiation, we observed deposition of CS lignin in “a string of pearl” manner in WT and *sgn3-3* (Fig. 1A). No endodermal lignification was observed in *myb36-2* or *sgn3-3 myb36-2* at this developmental stage of the root (Fig. 1A). Later in endodermal development, 10 cells after the onset of elongation, we observed a continuous CS ring of lignin, sealing the endodermal cells in WT plants (Fig. 1A-C). As we expected, in *sgn3-3* impaired in the activation of the Schengen-pathway, CS lignification still appears in a discontinuous fashion, and *myb36-2* exhibits compensatory lignification in the corners of the endodermal cells facing the cortical side of the endodermis as previously reported (9, 17). In contrast, no ectopic lignification was observed in the double mutant *sgn3-3 myb36-2* at the same developmental stage (Fig. 1A-C). These results establish that the cell-corner compensatory lignification observed in *myb36-2* lacking CS (17), is SGN3-dependent.

**Figure 1.**
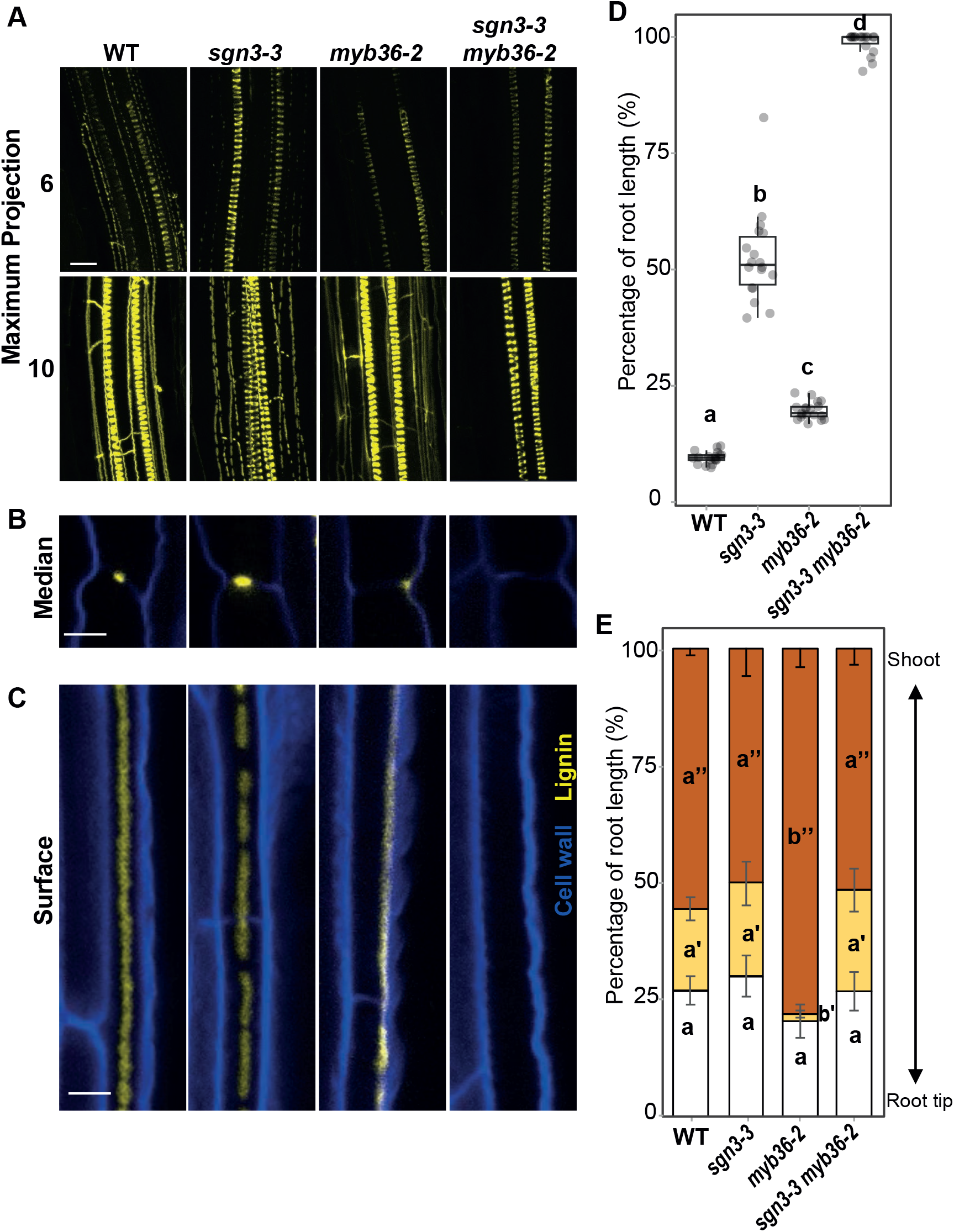
Disruption of *MYB36* and *SGN3* abolish endodermal lignification and root apoplastic barrier. **(A)** Maximum projection of lignin staining at the 6^th^ and 10^th^ endodermal cell after the onset of elongation. Spiral structures in the centre of the root are xylem. Scale bar = 10 μm. Median (B) and surface (**C**) view of an endodermal cell at 10 cells after the onset of elongation. Scale bar = 5 μm. The roots were cleared and stained with basic fuchsin (yellow) for lignin and with Calcofluor white (blue) for cellulose. (**D**) Boxplot showing the percentage of the root length permeable to propidium iodide. n=18 from two independent experiments. Different letters represent significant differences between genotypes using a Mann-Whitney test (p<0.01). (**E**) Quantification of suberin staining along the root. The results are expressed in percentage of root length divided in three zones: unsuberised (white), discontinuously suberised (yellow), continuously suberised (orange). n = 6, error bars: SD. Individual letters show significant differences using a Mann-Whitney test between the same zones (p<0.01).

To test how these different patterns of endodermal lignification found in WT, *sgn3-3*, *myb36-2* and *sgn3-3 myb36-2* affect the permeability of the root apoplast, we assessed the penetration of the fluorescent apoplastic tracer propidium iodide (PI) (23) into the stele (Fig. 1D). We quantified the percentage of root length permeable to PI, and found that it is partially increased in *sgn3-3* and *myb36-2* in comparison with WT (Fig. 1D). Surprisingly, we observed in the double mutant *sgn3-3 myb36-2*, that the entire length of the root was permeable to PI, indicating an additive effect of both mutations in the double mutant. This result suggests that MYB36 and SGN3 control endodermal lignification through two-independent pathways. The lack of compensatory cell-corner lignification in *sgn3-3 myb36-2* could explain the full permeability of the root found in these plants. This finding supports recent observations assigning a role as an apoplastic barrier to the SGN3-dependent cell-corner lignification (13). In addition, over-activation of the Schengen-pathway is also known to trigger an enhanced suberisation in other CS mutants, including *myb36* (9, 17, 24). We confirm this observation in the *myb36-2* mutant where an early suberisation is observed (Fig. 1E). This enhanced suberisation in *myb36* is also SGN3 dependent since the *sgn3-3 myb36-2* double mutant shows the same pattern of suberisation as WT plants (Fig 1E).

Our results indicate that MYB36 and SGN3 control endodermal lignification through two pathways: (a) The pathway involved in CS lignification controlled by both MYB36 and SGN3; and (b) The pathway involved in compensatory lignification of the endodermal cell corners controlled exclusively by SGN3.

### Endodermal cell-corner lignin is chemically distinct from CS lignin

We investigated the chemical nature and biochemical origins of CS lignin and compensatory cell-corner lignin. For this, we used confocal Raman microscopy on root cross-sections, in order to spatially resolve the chemistry of these different types of lignin. We triggered endodermal cell-corner lignin deposition by feeding WT plants with CIF2 peptide (+CIF2), the ligand of the SGN3 receptor, able to activate the Schengen-pathway. We separately imaged regions of interest (ROIs) containing CS lignin in WT plants, ROIs with endodermal cell-corner lignin in WT treated with CIF2 and ROIs containing xylem lignin from WT plants, treated or not with CIF2 (Sup. Fig. 1). Then, we used a multivariate curve resolution (MCR) analysis on these Raman images to spatially and spectrally resolve lignin in these different ROI. The lignin spectra corresponding to these regions are shown in Fig. 2A-B. We observed that the CS lignin spectrum is distinct from that of endodermal cell-corner lignin. For example, peaks known to be assigned to lignin display higher (ex: 1337 cm^−1^, aliphatic OH bend (25)) and lower (ex: 1606 cm^−1^, aromatic ring stretch (25)) intensity in CS lignin in comparison with endodermal cell-corner lignin. Another striking difference was observed for the peak at 1656-1659 cm^−1^ assigned to a double bond conjugated to an aromatic ring (*e.g.*: coniferyl alcohol or coniferaldehyde, (25)). This peak is missing in the endodermal cell-corner lignin of WT treated with CIF2 in comparison with CS lignin, suggesting a change in the phenolic composition of the cell-corner lignin. Conversely, the xylem lignin spectrum of plants treated with or without CIF2 was similar, with the most intense peaks showing comparable intensity. This suggests that changes in lignin composition triggered by the over-activation of the Schengen-pathway mainly occur in the endodermis, and xylem lignin remains largely unaffected.

**Figure 2.**
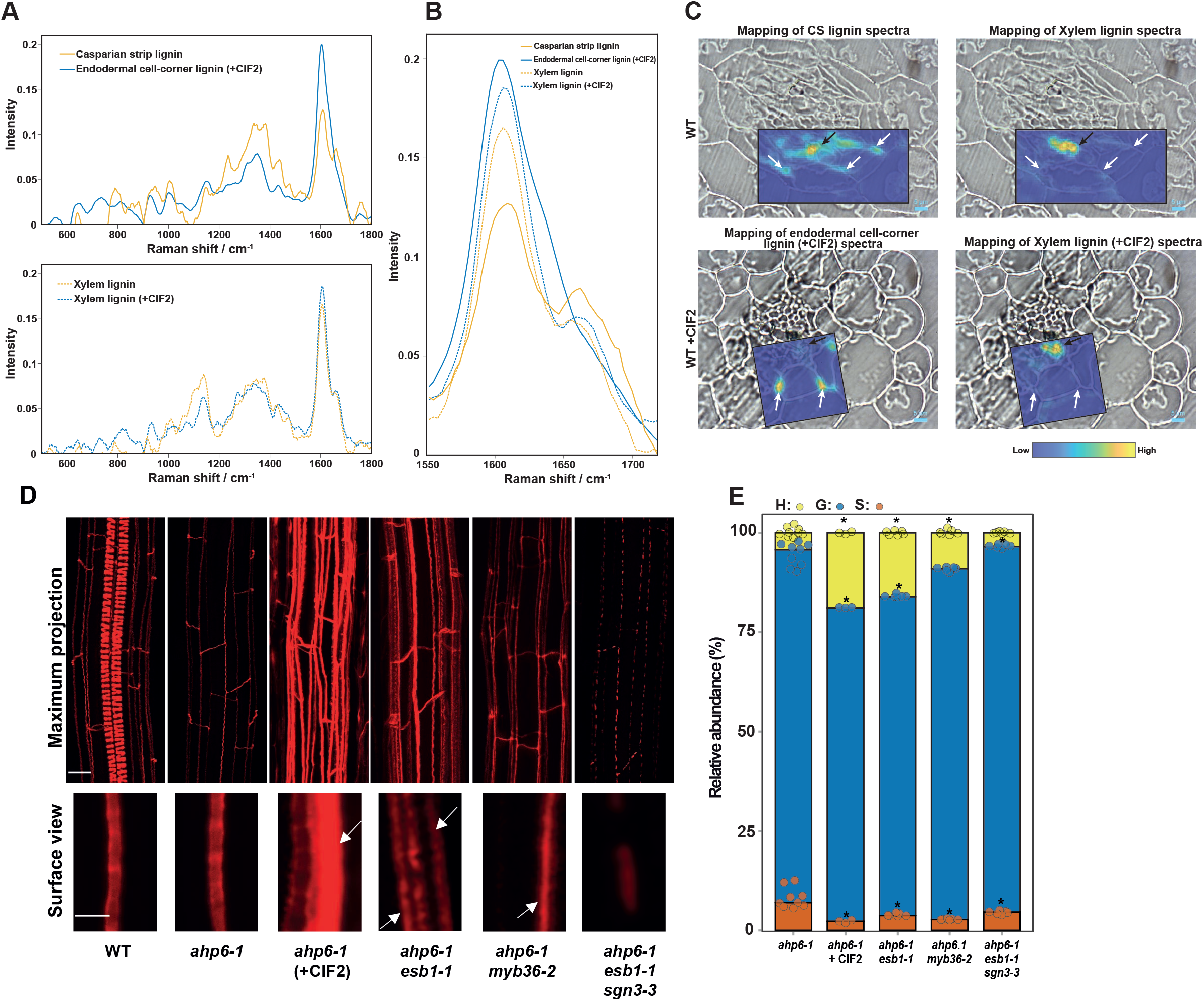
Activation of the Schengen-pathway triggers the deposition of a distinct “stress” lignin in the endodermis. (**A**) Raman spectra of lignin of the different regions of interest presented in Sup. Fig. 1 and determined using a Multivariate Curve Resolution (MCR) analysis. The MCR analysis was performed on small Raman maps from independent plants containing CS lignin of WT (n = 8), cell-corner lignin of WT treated with CIF2 (+CIF2; n = 5) and for xylem lignin of WT (n = 2) and xylem lignin of WT treated with CIF2 (n = 2). (**B**) Close view of Raman spectra presented in (A) in the lignin aromatic region between 1550cm^−1^ and 1700 cm^−1^. (**C**) Large Raman maps of roots of WT and WT treated with CIF2 (+CIF2). The intensity of the different lignin spectra presented in (A) was mapped onto large Raman maps containing xylem and endodermal lignin. (**D**) Lignin staining with basic fuchsin at a distance of 3 mm from the root tip in WT, *ahp6-1*, *ahp6-1esb1-1*, *ahp6-1esb1-1sgn3-3*, *ahp6-1myb36-2* and *ahp6-1* treated with CIF2. The plants were grown for 6 days in presence of 10 nM 6-Benzylaminopurine (BA). Upper panel shows a maximum projection of the root (Scale bar = 10 μm). Spiral structures in the centre only observed in the WT root are protoxylem. Lower panel shows surface view of endodermal cells (Scale bar = 5 μm). White arrows indicate ectopic lignification. (**E**) Relative abundance of the lignin monomers released by thioacidolysis (*p*-hydroxyphenyl (H), guaiacyl (G), and syringyl (S) units) in root tips of *ahp6-1* (n=9), *ahp6-1* treated with CIF2 (+CIF2; n = 3), *ahp6-1esb1-1* n = 6, *ahp6-1 myb36-2* (n = 6) and *ahp6-1 esb1-1 sgn3-3* (n = 6). Asterisks represent significant differences from the *ahp6-1* control for each individual monomer using a Mann-Whitney U test (*p*-value < 0.01).

These conclusions were further confirmed spatially by mapping the intensity of these different lignin spectra on large Raman maps containing xylem and endodermal lignin in WT plants treated or not treated with CIF2 (Fig. 2C). We observed that the CS lignin spectrum localises to the CS and xylem vessels suggesting a similar lignin composition, as previously shown for monomer composition using thioacidolysis (7). Additionally, the endodermal cell-corner lignin spectrum localises almost exclusively to the site of lignification in the corners of the endodermal cells, and is essentially absent from the xylem. However, the xylem lignin spectrum (WT +CIF2) matches exclusively to the xylem vessel and is not observed at the endodermal cell corners. This strongly supports the conclusion that over-activation of the Schengen-pathway triggers deposition of lignin at endodermal cell-corners that has a unique chemical composition compared to both CS and xylem lignin.

To confirm these differences between CS and endodermal cell-corners lignin, we adopted an approach to directly measure the subunit composition of endodermal lignin avoiding possible contamination from the highly lignified protoxylem cells (7). We genetically crossed a collection of CS mutants that represent different level of lignin accumulation in the endodermis with the *arabidopsis histidine transfer protein 6.1* mutant (*ahp6-1*). This mutant, in the presence of low amounts of the phytohormone cytokinin, shows a strong delay in protoxylem differentiation, without affecting CS formation (Fig. 2D) (7, 26). Therefore, in the resulting lines the majority of lignin derived from the protoxylem is lost allowing us to analyse primarily lignin with an endodermal origin. To explore how the chemical composition of the cell-corner lignin differs from CS lignin, we collected root tips (3 mm) of 6-day-old *ahp6-1* and *ahp6-1 esb1-1 sgn3-3* mutants accumulating CS lignin only, and from mutants (*ahp6-1 myb36-2* and *ahp6-1 esb1-1*) with cell-corner lignification and a reduced amount of CS lignin. Additionally, as a control we used *ahp6-1* plants treated with the CIF2 peptide that strongly induces the Schengen-pathway and deposition of cell-corner lignin (Fig. 2D). We measured the relative content of H, G and S subunits in lignin in all samples using thioacidolysis followed by GC-MS (Fig. 2E). We found that CS lignin monomer composition in our control line *ahp6-1* (H: 5%, G: 87%, S: 8%) was similar to that previously reported (7). The monomer composition of the defective CS in the mutant *ahp6-1 esb1-1 sgn3-3* is overall similar to WT with a small increase in G and decrease in S subunits. Strikingly, we observed that lignin composition in the lines and treatments that induce the accumulation of cell-corner lignin (*ahp6-1 esb1-1, ahp6-1 myb36-2, ahp6-1*(+CIF2)) was different from the control and mutant lines that only accumulate lignin in the CS. The lignin extracted from these plants showed a higher proportion of H monomers. In the case of *ahp6-1* treated with CIF2, H content was increased to 19 % and G content was decreased. Thioacidolysis and Raman results indicate that over-activation of the Schengen-pathway triggers the deposition of a chemically distinct H-rich lignin in the corner of the endodermal cells.

Such a high content of H subunits in lignin is rarely found in angiosperm. Similar levels of H subunits in lignin mainly occurs in compression wood of gymnosperm (27–30) and in defence-induced lignin, and has been termed “stress lignin” (31–35). We therefore conclude that Schengen-pathway induced endodermal cell-corner lignin is a novel form of ‘stress lignin’. Taken together, both chemical analysis of lignin subunits by thioacidolysis and spatially resolved confocal Raman spectroscopy show that lignin deposited in endodermal cell corners upon activation of the Schengen-pathway is H-rich and chemically and spatially distinct from both CS and xylem lignin.

### Schengen-pathway modulates the phenylpropanoid pathway and induces defense-related mechanisms

To investigate the biosynthesis of the endodermal H-rich stress lignin we performed RNA-seq on root tips (5 mm) of WT plants, on roots showing a strong activation of the Schengen-pathway (WT treated with exogenous CIF2, *myb36-2* and *esb1-1*) and roots with no Schengen signalling (*sgn3-3*, *esb1-1 sgn3-3*, *sgn3-3 myb36-2* and *sgn3-3* treated with exogenous CIF2). Clustering analysis of the differentially expressed genes shows that roots displaying cell-corner lignification (WT treated with exogenous CIF2, *myb36-2* and *esb1-1*) due to the over-activation of the Schengen-pathway share a similar transcriptional response that is distinct from that observed in the other genotypes (Fig. 3A, Sup. Fig. 2A, Sup. Table 1). CIF2 application to *sgn3-3* shows a similar transcriptional response to non-treated WT and *sgn3-3* and does not trigger the transcriptional changes observed during the strong activation of the Schengen-pathway (WT treated with exogenous CIF2, *myb36-2* and *esb1-1*). This is in line with previous transcriptomic data in response to CIF2 (18) and the idea that SGN3 is the only receptor for CIF2 in roots. We observed that genes in cluster C1 are upregulated by the activation of the Schengen-pathway. This cluster is enriched in genes involved in the phenylpropanoid pathway (Sup. Fig. 2B). We hypothesize that the activation of this pathway would provide the phenolic substrates required for the enhanced lignification and suberisation induced by the Schengen-pathway. We observed strong activation of expression of genes encoding all the key enzymes of the phenylpropanoid pathway required for monolignol biosynthesis, with the exception of *C3’H*, *C3H* and *F5H* (Fig. 3B, Sup. Fig. 2C). H-rich lignin is known to be accumulated when expression of *C3’H* is repressed in *A. thaliana* and poplar (36–39). This activation of all the main enzymes of the phenylpropanoid pathway with the exception of *C3’H* observed after triggering the Schengen-pathway could explain the high level of H-units incorporation into endodermal cell-corner lignin (Fig. 3B, Sup. Fig. 2C). Similarly, the roots of the cellulose synthase isomer mutant *eli1* (*ectopic lignification1*) accumulate H-rich lignin and display strong gene activation for most of the phenylpropanoid pathway, with the exception of *C3’H* (40). Interestingly, ectopic lignification in this mutant is also under the control of another receptor-like kinase, *THE1* (THESEUS), also involved in cell wall integrity sensing (41, 42). We then tried to identify transcriptional regulators with a role in the regulation of the Schengen-pathway sector controlling phenylpropanoid synthesis. We performed a gene expression correlation analysis between the phenylpropanoid pathway genes and their transcriptional regulators (3, 43) (Sup. Fig. 2C). We found that the expression of the transcription factor *MYB15* highly correlates with the expression of most of the genes required for monolignol biosynthesis, with the notable exception of *C3’H* (Sup. Fig. 2C). Upregulation of *MYB15* in response to CIF2 have been previously shown (18). This transcription factor is known to bind to the promoter of *PAL1*, *C4H*, *HCT*, *CCoAOMT1* and *COMT* but does not bind to the promoter of *C3’H* and *F5H* (44). Schengen-pathway activation of *MYB15* expression provides a plausible mechanism to explain the induction of the main enzymes of the phenylpropanoid pathway with the exception of *C3’H* and *F5H*. This modulation of gene expression could explain the enhanced incorporation of *p*-coumaryl alcohol into the stress lignin we observe at endodermal cell corners. Interestingly, *MYB15* is an activator of basal immunity in *A. thaliana* by inducing the synthesis of defense lignin and soluble phenolics (44).

**Figure 3.**
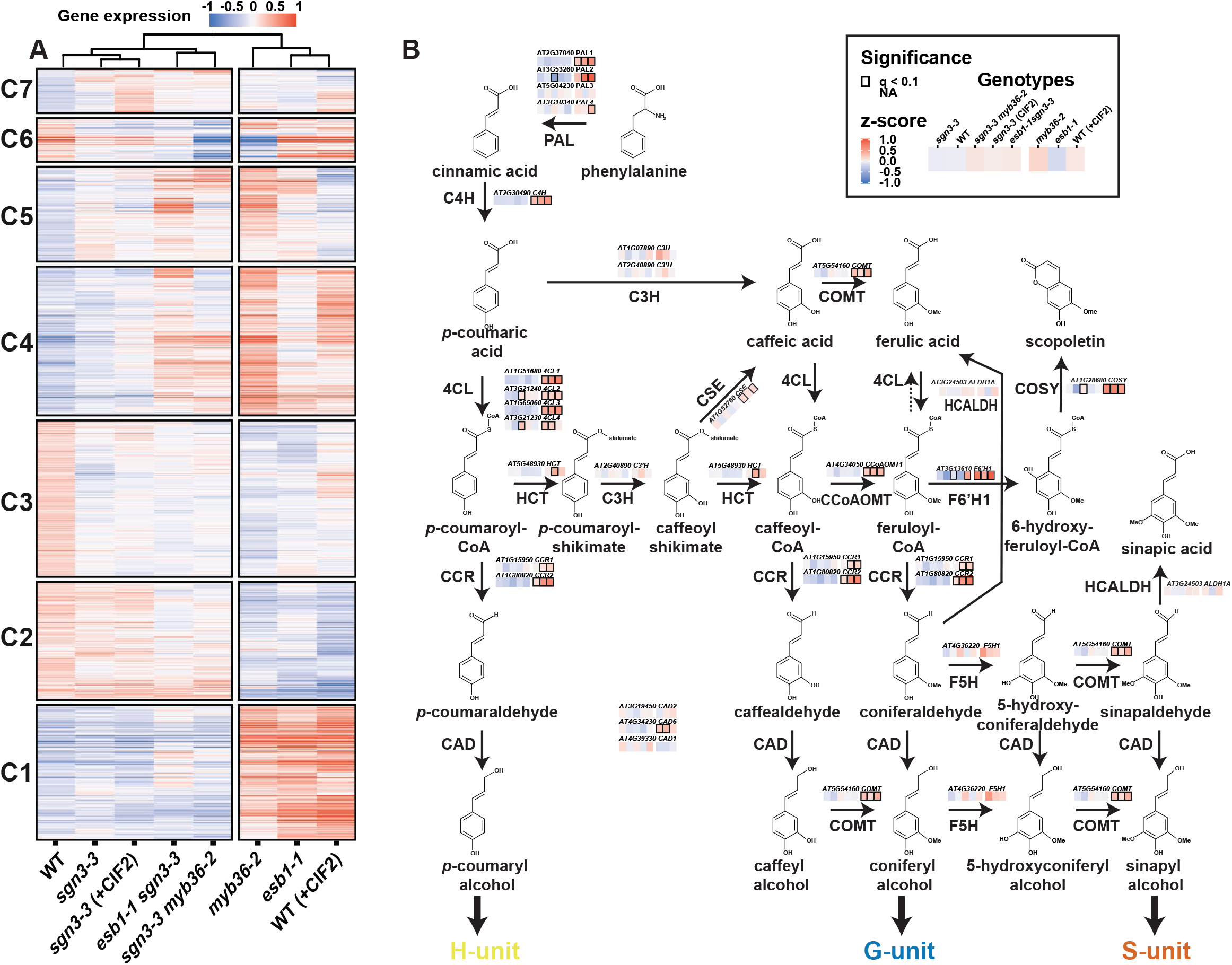
Modulation of the phenylpropanoid pathway by the Schengen-pathway. (**A**) Heatmap of the 3266 differentially expressed genes identified in the RNA-seq in root tips of wild-type (WT), *sgn3-3*, *esb1-1*, *myb36-2*, *esb1-1 sgn3-3*, *sgn3-3 myb36-2* plants. Treatment with 100 nM CIF2 was applied as indicated (+CIF2) for WT and *sgn3-3* plants. Clusters (C) are designated with numbers (n = 6). Genes belonging to each cluster are listed in Sup. Table 1. (**B)** Phenylpropanoid pathway leading to the lignin monomers and scopoletin biosynthesis (adapted from and (68)). Solid arrows represent enzymatic steps. Gene expression from the genes selected in Sup. Fig. 3C were mapped on the pathway according to their KEGG enzyme nomenclature. Only the genes with a demonstrated function in lignin biosynthesis as listed in Sup. Fig. 3C were mapped. PAL, PHENYLALANINE AMMONIA-LYASE; C4H, CINNAMATE 4-HYDROXYLASE; 4CL, 4-COUMARATE:CoA LIGASE; HCT, *p*-HYDROXYCINNAMOYL-CoA:QUINATE/SHIKIMATE *p*-HYDROXYCINNAMOYLTRANSFERASE; C’3H, *p*-COUMARATE 3’-HYDROXYLASE; C3H, COUMARATE 3-HYDROXYLASE; CSE, CAFFEOYL SHIKIMATE ESTERASE; CCoAOMT, CAFFEOYL-CoA O-METHYLTRANSFERASE; CCR, CINNAMOYL-CoA REDUCTASE; F5H, FERULATE 5-HYDROXYLASE; COMT, CAFFEIC ACID O-METHYLTRANSFERASE; CAD, CINNAMYL ALCOHOL DEHYDROGENASE; HCALDH, HYDROXYCINNAMALDEHYDE DEHYDROGENASE; COSY, COUMARIN SYNTHASE; F6’H1 FERULOYL COA ORTHO-HYDROXYLASE 1.

To test if over-activation of the Schengen-pathway leads to the production of defense-inducible soluble phenolics, we undertook secondary metabolites profiling using Ultra High Performance Liquid Chromatography (UHPLC). Profiling was performed on root tips (5 mm) of the *esb1-1* mutant having a defective CS and constitutive activation of the Schengen-pathway, in *sgn3-3* and *sgn3-3 esb1-1* having a defective CS and inactivation of the Schengen-pathway and in WT. We observed distinct accumulation of soluble secondary metabolites across the different genotypes (Sup. Fig. 3, Sup. Table 2). We identified 20 phenolic compounds that differentially accumulate specifically due to the activation of the Schengen-pathway out of 52 compounds. We found higher accumulation of the conjugated neolignan G(8-O-4)*p*CA, scopoletin, flavonoid derivatives such as conjugated kaempferol (astragalin and 4′-O-acetylkaemferol-3-O-hexoside), isorhamnetin and acetylhyperoside. Scopoletin biosynthesis is controlled by the enzyme F6’H1 and COSY (45, 46) and the transcription factor MYB15 (44). We found that the expression of the three genes encoding these proteins is induced by the over-activation of the Schengen-pathway (Fig 3B, Sup. Fig. 2C). Scopoletin is a modulator of plant-microbe interaction (44, 47–49). In addition to that, we found a strong induction of genes related to defense (response to chitin/systemic acquired resistance/immune response/hypersensitive response) among the genes induced by the activation of the Schengen-pathway (C1; Fig. 3A and Sup. Fig. 2B). This corroborates a publication showing similarities between the Schengen-pathway and the microbe-associated molecular patterns (MAMP) signalling pathway (18).

### Cell wall attachment to plasma membrane relies on CS domain formation rather than lignin deposition

The apoplast in between two endodermal cells is sealed by the deposition of CS lignin. This sealing is perfected by the anchoring of the CS membrane domain (CSD) to the cell wall (CW), through an unknown mechanism. Upon plasmolysis, the protoplasts of endodermal cells retract but the CSD remains tightly attached to the CS (23, 50, 51). This attachment appears in a developmental manner during the differentiation of the endodermis and occurs in a concomitant manner with the recruitment of the Casparian strip membrane domain proteins (CASPs) at the CSD and with CS lignin deposition (23). We then wanted to study whether or not the different types and sites of lignification contribute to the attachment of the plasma membrane (PM), to the CW. To visualize the PM, we used an endodermis specific PM marker (pELTP::mCit-SYP122) in WT, *sgn3-3*, *myb36-2* and *sgn3-3 myb36-2* (Fig. 4A). The PM marker is excluded from the CSD in WT as described for other endodermal plasma membrane marker lines (14, 23). This exclusion is still observed in *sgn3-3* but in an interrupted manner similarly to that observed for lignin (Fig. 1A-C). The exclusion domain disappears entirely in the *myb36-2* and *sgn3-3 myb36-2* mutants. Additionally, no exclusion zone in the PM is observed in *myb36-2* where cell-corner lignin is deposited. We then used mannitol-induced plasmolysis to visualise the PM attachment to the CW. Upon plasmolysis, the PM retracts but remains attached to the CS in WT and *sgn3-3*, forming a flattened protoplast (Fig. 4A). However, small portions of the PM are able to detach from the CW in a *sgn3-3* mutant as seen in Supp. Fig. 4A. This is likely to happen where the PM exclusion domain is interrupted in *sgn3-3* (Fig. 4A). In *myb36-2* and *sgn3-3 myb36-2*, the CW attachment to the PM is lost (Fig. 4A, Supp. Fig. 4A). Importantly, retraction of the PM is observed in *myb36-2* at the corner of the endodermal cells on the cortex side where cell-corner lignin is deposited (Fig 1A). These results clearly show the requirement of MYB36 for the formation of the CSD excluding the PM marker. Additionally, the presence of CSD, but not cell-corner lignin is required for PM attachment to the CW.

**Figure 4.**
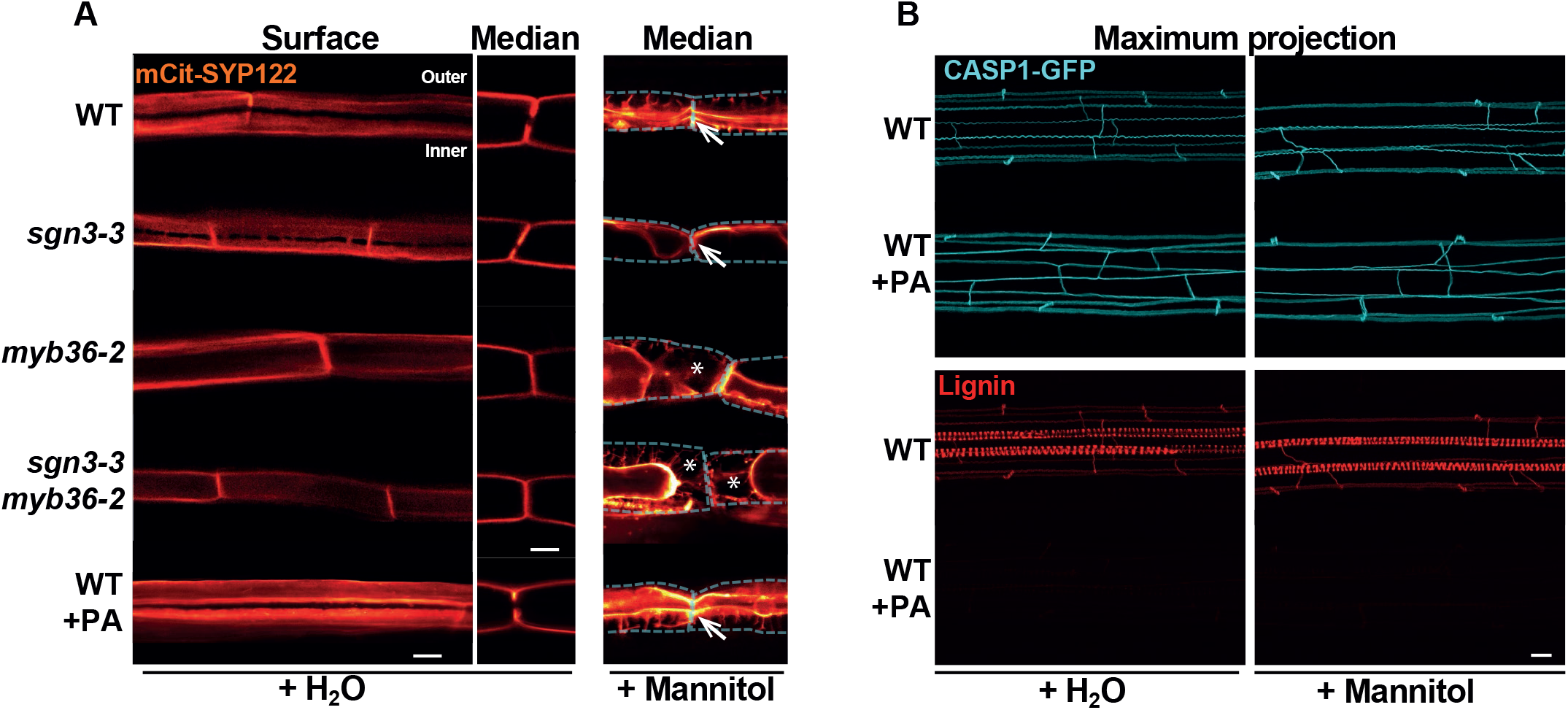
PM attachment to the CW is MYB36-dependent but does not rely on lignin deposition. (**A**) Median and surface view of the endodermal plasma membrane using the marker line pELTP::SYP122mCitrine before plasmolysis (+H2O) and after plasmolysis (+Mannitol) at 15 cells after the onset of elongation. WT plants were treated or not from germination with 10 μM piperonylic acid (+PA). White asterisks show the exclusion domain at the CSD. The dashed line represents the contours of the cells before plasmolysis. Arrows show the plasma membrane attachment to the cell wall. Blue asterisks show the plasmolysis generated space where no attachment is observed. Scale bar = 5 μm. “inner” designates the stele-facing endodermal surface, “outer”, the cortex-facing surface. (**B**) Maximum projection of CASP1-GFP and lignin staining with basic fuchsin in cleared roots from plants grown with or without 10μM piperonylic acid and subjected to plasmolysis with Mannitol. Scale bar = 10 μm.

We then tested if CS lignin is required for the PM attachment to the CS. For this, we used an inhibitor of the phenylpropanoid pathway, piperonylic acid (PA), that inhibit lignin accumulation (7). Treatment with PA suppresses lignin accumulation in the vasculature and in the CS (Fig. 4B). Absence of lignin did not affect the exclusion of the PM marker at the CSD. This was further confirmed using the CSD marker line (pCASP1::CASP1-GFP) (12) that showed similar localisation independently of the CS lignin presence (Fig. 4B). Additionally, the PM attachment to the CS is still observed when CS lignin deposition is inhibited (Fig. 4A-B, Supp. Fig. 4A). These findings indicate that CS lignin is not required for the formation of the CSD as previously reported (14, 18) and importantly that CS lignin does not participate in anchoring the CSD to the CW. Other CW compounds might be involved in that process.

The absence of PM attachment to the site of cell-corner lignification is likely to affect the permeability of the apoplast of the endodermal cells. This can consequently affect the transport of water and solutes to the shoot.

### Total absence of an endodermal apoplastic barrier triggers major ionomic changes

We then investigated how the different types of endodermal lignification control nutrient homeostasis in the plant. The *sgn3-3* (delayed CS barrier, no cell-corner lignin), *myb36-2* (no CS lignin, has cell-corner lignin) and *sgn3-3 myb36-2* (no CS or cell-corner lignin) mutants were grown using different growth conditions (agar plate, hydroponic, and natural soil) and leaves were analysed for their elemental composition (ionome) using inductively coupled-mass spectrometry (ICP-MS; Fig. 5A and Sup. Table 3). A Principal Component Analysis (PCA) of the ionome of leaves reveals that all the mutants have different leaf ionomes compared to WT when grown on plates (Fig. 5B), in hydroponic and to a lesser extent in natural soil (Sup. Fig. 5 A-B). Based on the PC1 axis, the double mutant *sgn3-3 myb36-2* displayed the most distinctionomic phenotype (Fig 5B, Sup. Fig 5 A-B). In line with our previous results (Fig. 1D), this effect indicates an additivity of the two mutations on the leaf ionome. Importantly, this result also supports that cell-corner stress lignin in the single mutant *myb36* can act as an apoplastic barrier to mineral nutrients.

**Figure 5.**
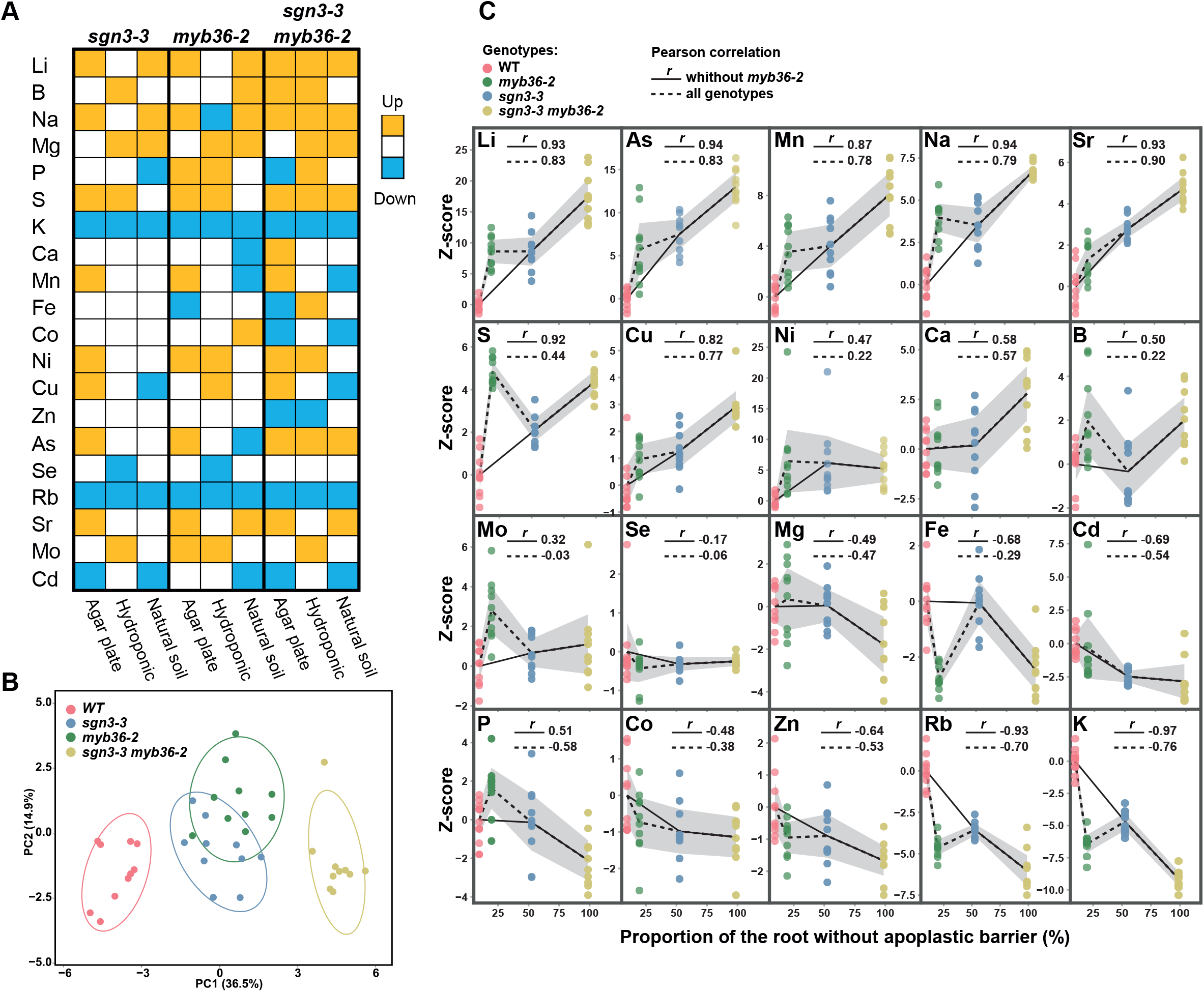
Absence of endodermal apoplastic barrier triggers major ionomic changes. **(A)** Overview of ions accumulation in shoot of *sgn3-3*, *myb36-2* and *sgn3-3 myb36-2* mutants compared to WT using different growth conditions in agar plates (long day, n=10), in hydroponics (short day, n=6) and natural soil (short day, n≥13). Elements concentration were determined by ICP-MS and are available in the Sup. Table 3. Colour code indicates significant changes in accumulation compared with the WT using a *t* test (p<0.01). **(B)**Principal component analysis (PCA) based on the concentration of 20 elements in shoots of plants grown in agar plates. Ellipses show confidence level at a rate of 90%. n=10 (**C**) Plots presenting the correlation between the z-scores of elements content in shoots of plants grown in agar plates of WT, *myb36-2*, *sgn3-3* and *sgn3-3 myb36-2* against the portion of root length permeable to propidium iodide as determined in Fig. 1D. The black lines show the average and the grey area show the 95% confidence interval (n = 10).

We next tested the correlation between the gradient of root apoplastic permeability across WT, *myb36-2*, *sgn3-3* and *sgn3-3 myb36-2* determined in Fig. 1D with their elemental content in leaves (Fig 5C). We observed that the *myb36-2* mutant does not fit into this correlation analysis as well as the other genotypes. This is likely due to the activation of the Schengen-pathway leading to the deposition of endodermal cell-corner stress-lignin, early suberisation, reduced root hydraulic conductivity, activation of ABA signalling in the shoot, and stomata closure (9, 24). Additionally, the *myb36-2* mutation interferes with overall root development (Sup. Fig. 5 C-E) as previously reported (52). This is due to the over-activation of the Schengen-pathway as the double *sgn3-3 myb36-2* mutant shows normal root development. Removal of *myb36-2* from the correlation analysis, leaving just lines with an inactive Schengen-pathway, improved the Pearson correlation coefficient for almost all the elements, and we observed a strong correlation (*r* ≥ 0.5 or *r* ≤ −0.5) for 15 out of 20 elements. We observed a strong positive correlation between an increased CS permeability and leaf accumulation of lithium (Li), arsenic (As), manganese (Mn), sodium (Na), strontium (Sr), sulphur (S), copper (Cu), calcium (Ca), boron (B), and a strong negative correlation with iron (Fe), cadmium (Cd), phosphorus (P), zinc (Zn), rubidium (Rb) and potassium (K). This suggests that a functional apoplastic barrier is required to limit the loss of essential elements such as K, Zn, Fe and P. Conversely, a defective apoplastic barrier allows increased leaf accumulation of the essential nutrients Mn, S, Cu, Ca, and B. These gradients of higher and lower accumulations of mineral nutrients and trace elements illustrate the bidirectional nature of the CS barrier, by blocking some solutes from entering the vasculature and by facilitating the accumulation of other solutes in the stele for translocation.

### Root hydraulic conductivity is reduced by the activation of the Schengen-pathway and it is not affected by the absence of endodermal lignification

We then measured the capacity of the root to transport water, also called root hydraulic conductivity (Lpr), in 3-week-old plants grown hydroponically. We observed that the root hydraulic conductivity remains unchanged in the mutants *sgn3-3* and *sgn3-3 myb36-2* in comparison with WT (Fig. 6A). In contrast, the *myb36-2* mutant showed a strong reduction of root hydraulic conductivity. These results established that the CS-based endodermal apoplastic seal does not control root water transport capacity, as in the absence of any barriers in *sgn3-3 myb36-2* (Fig. 1D) root hydraulic conductivity is the same as WT. This is consistent with water transport occurring mainly via the transcellular pathway, with a major contribution via aquaporins (53). The reduced hydraulic conductivity observed in *myb36-2* is consistent with that previously observed in *esb1* which also has an activated Schengen-pathway (24). The reduced hydraulic conductivity in *esb1* originates mainly from a reduction in aquaporin-mediated water transport as determined using an pharmacological approach (24). Here, our RNA-seq experiment revealed a GO-term enrichment in cluster C2 (genes repressed by the Schengen-pathway, Fig. 3A) relating to water deprivation (Sup Fig 3B) and importantly, 10 aquaporin genes are down regulated by activation of the Schengen-pathway (Fig 6B). This set of aquaporin genes contains several highly expressed aquaporins in root, including *PIP2,2* known to significantly contribute to root hydraulic conductivity (54, 55). This would provide an explanation for the reduction in root hydraulic conductivity observed in both *myb36* and *esb1*.

**Figure 6.**
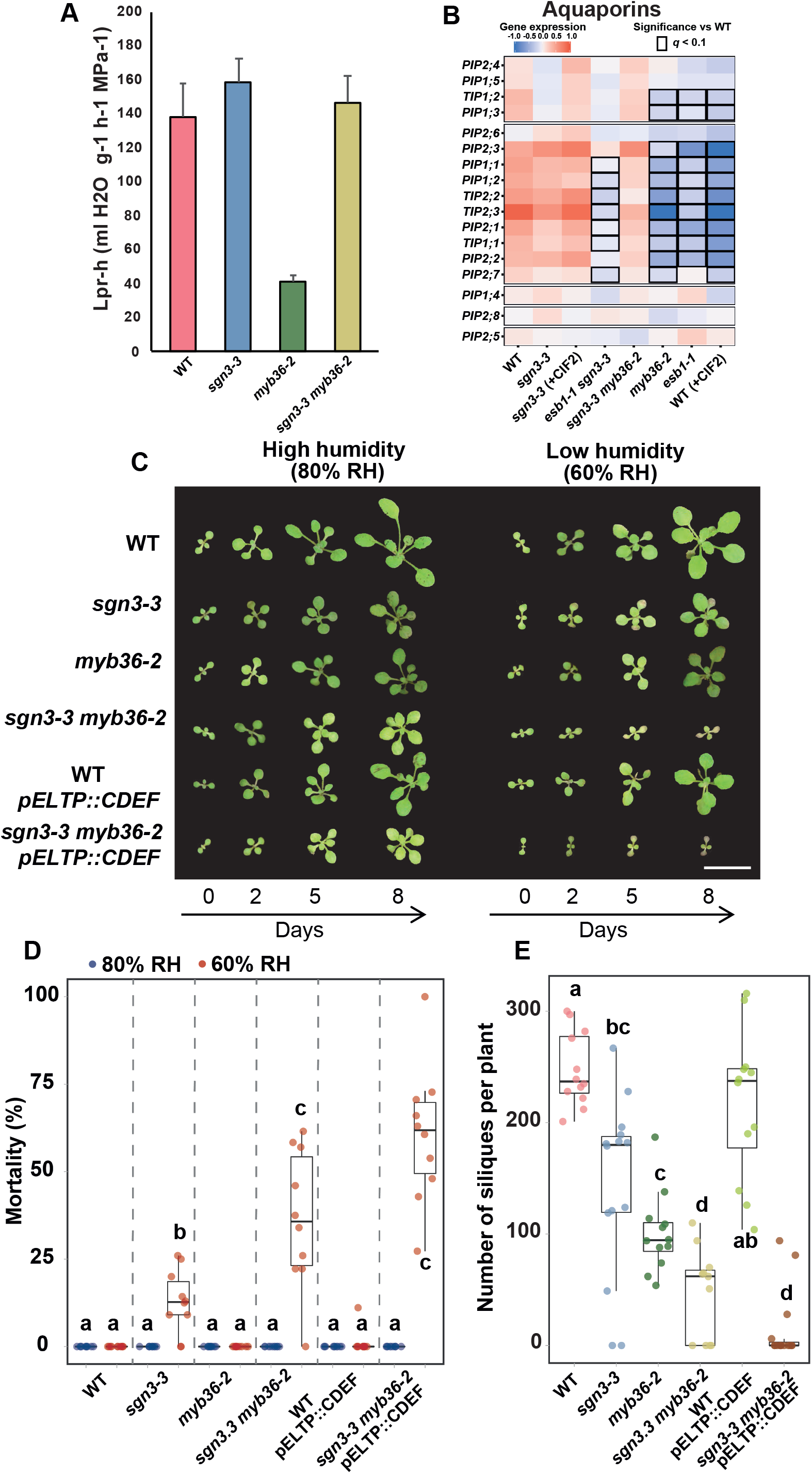
Activation of the Schengen-pathway represses water transport and maintains plant growth, survival and fitness under fluctuating environment. (**A**) Hydrostatic root hydraulic conductivity (*L*pr-h) in WT, *sgn3-3*, *myb36-2*, *sgn3-3 myb36-2* grown hydroponically for 19-21 days under environmental controlled conditions. Hydraulic conductivity was measured using pressure chambers (*L*pr-h) (means ± SE, n ≥ 3). (**B**) Heatmap of aquaporins expression across the different genotypes and treatments used in the RNAseq experiment. **(C)**Representative pictures of WT, *sgn3-3*, *myb36-2*, *sgn3-3 myb36-2,* WT - *pELTP::CDEF* and *sgn3-3 myb36-2* - *pELTP::CDEF* plants germinated in soil with a high humidity (80%) for 7 days and then transferred in an environment with a lower (60% RH) or with constant humidity (80% RH). Pictures were taken 0, 2, 5 and 8 days after the transfer. Scale bar = 1 cm. (**D)**Boxplots showing the proportion of dead plants after transfer in an environment with constant humidity (80% RH, blue) or with a lower (60% RH, red). The plants displaying no growth after 9 days and showing necrosis in all the leave surface were considered as dead plants. Each point represents the proportion of dead plants in a cultivated pot compared to the total number of plants for one genotype in the same pot. Pots were containing at least 8 plants of each genotypes, n=10 pots. Different letters represent significant differences between genotypes using a Mann-Whitney test (p<0.01). **(E)**Boxplots showing the number of siliques produced per plants. Plants were cultivated in a high humidity environment for 10 days after germination and then transferred to a greenhouse. Each point represents the total number of seeds containing siliques per plant (n≥12). Different letters represent significant differences between genotypes using a Mann-Whitney test (p<0.01).

### Endodermal lignification is required for plant growth and survival in low humidity

Given the significant impacts that CS and Schengen-pathway activation have on mineral nutrient homeostasis (Fig. 5) and water transport (Fig. 6A-B), we further investigated their impact on growth and development. The double mutant *sgn3-3 myb36-2* displayed a severe dwarf phenotype when grown hydroponically or in natural soil but not on agar plates in comparison with WT and the single mutants (Supp. Fig. 5 C-G). This indicates a critical role of CS for maintaining normal plant growth and development. However, this is conditioned by the growth environment. The high humidity environment and consequently reduced transpiration of plants on agar plates in comparison with the other growth environments could explain these phenotypical differences. Indeed, reduced leaf transpiration is a key part of the compensation mechanisms mitigating the loss of CS integrity allowing relatively normal growth as reported in (24). We then tested if differences in relative humidity (RH) can affect plant growth in the absence of an endodermal root barrier. For this, we used *sgn3-3* (delayed CS barrier, no cell-corner lignin), *myb36-2* (no CS lignin, has cell-corner lignin) and *sgn3-3 myb36-2* (no CS or cell-corner lignin). Additionally, we tested if the presence of endodermal suberin can affect plant growth by using lines expressing the Cutinase DEstruction Factor (CDEF) under the control of an endodermis specific promoter (pELTP::CDEF) in a WT and *sgn3-3 myb36-2* background and showing a strong reduction of endodermal suberin deposition (Supp. Fig. 6A). Seedlings were germinated in soil in a high humidity environment (80% RH) for 7 days and transferred to an environment with the same (80% RH) or lower (60% RH) humidity. We measured the leaf surface area as a proxy of plant growth (56) at 0, 2, 5 and 8 days after transfer (Fig. 6C and Supp. Fig. 6B). In a high and low humidity environment, all mutants with reduced CS functionality (*sgn3-3*, *myb36-2*, *sgn3-3 myb36-2* and *sgn3-3 myb36-2-pELTP::CDEF*) show a reduction of leaf surface in comparison with WT. Importantly, the growth reduction observed in the absence of endodermal lignification (*sgn3-3 myb36-2*) is severe, specifically in low humidity conditions, in comparison with all other genotypes and high humidity conditions. The *sgn3-3 myb36-2* plants with no growth after 9 days started to display necrosis over all the leave surface and were considered dead as quantified in Fig. 6D. Low humidity triggers a high percentage of mortality in *sgn3-3 myb36-2* and to a lesser extent in *sgn3-3* compared to WT and to the other genotypes in which no mortality is observed when grown in low humidity conditions. Such mortality was not observed if *sgn3-3 myb36-2* was grown in high humidity. This highlights that endodermal lignification is required for maintaining plant growth and survival under low humidity. However, this is not the case for endodermal suberisation because the removal of suberin by expressing CDEF in WT and *sgn3-3 myb36-2* did not affect mortality and leaf surface area after 8 days at a lower humidity in comparison with their respective backgrounds (Fig.6 C,D, Supp. Fig. 6B).

The strong growth reduction observed in *sgn3-3 myb36-2* in comparison with WT and the single mutants could be associated with the lack of root selectivity leading to major ionomic changes as shown in Fig. 5. The low humidity would generate a higher transpiration stream and consequently leads to a more uncontrolled and detrimental accumulation of mineral nutrient and trace elements in the leaves in comparison with high humidity. Alternatively, a high humidity environment would slow the transpiration rate, allowing plants to better control nutrient acquisition (24).

We then measured the impact of the absence of CS, suberin or of the constitutive activation of the Schengen-pathway on plant fitness. For this, we determined the number of seed-containing siliques per plant as an estimation of fitness (Fig. 6E). A significant reduction of silique numbers is observed in all the genotypes in comparison to WT with the exception of *pELTP::CDEF*. The mutant *sgn3-3* displaying partial root apoplastic barrier defects showed a decrease in siliques number in comparison with WT. A similar decrease was observed for *myb36-2* displaying also a partial root apoplastic barrier defect with cell-corner stress lignin deposition. Complete disruption of endodermal lignification strongly affects silique production as observed for *sgn3-3 myb36-2* and *sgn3-3 myb36-2 - pELTP::CDEF*. These results establish that the CS is essential for plant fitness. Further, activation of the Schengen-pathway helps protect the plant from this detrimental impact on fitness when the barrier function of the CS is compromised. The *sgn3-3 myb36-2* - *pELTP::CDEF* line reported here, with it complete lack of endodermal lignin and suberin extracellular barriers and Schengen-dependent signaling, is a powerful tool for studying the role of endodermal barriers in a range of processes such as nutrient, hormone and water transport and biotic interaction with soil microorganisms.

The data presented here revealed that the Schengen-pathway is involved in the deposition of two chemically distinct types of lignin. The Schengen-pathway with MYB36 are required for the deposition of CS lignin. Constitutive activation of the Schengen-pathway leads to the deposition of a chemically distinct stress-like type of lignin. This deposition of stress-lignin contributes to sealing the apoplast maintaining ion homeostasis in the absence of CS integrity. However, no PM attachment to the CW is observed at the site of stress-lignin deposition as seen for the CS, suggesting an inferior seal is formed.

## MATERIALS & METHODS

### Plant material

*Arabidopsis thaliana* accession Columbia-0 (Col-0) was used for this study. The following mutants and transgenic lines were used in this study: *sgn3-3* (SALK_043282) (9), *myb36-2* (GK-543B11) (17), pCASP1::CASP1-GFP (12), *ahp6-1* (26), *esb1-1* (10), *pELTP::CDEF*(57), pELTP::SYP122-mCitrine.

The corresponding gene numbers are: *SGN3*, At4g20140; *MYB36*, At5g57620; *CASP1*, At2g36100; *AHP6*, At1g80100; *ESB1*, At2g28670; *ELTP*, At2g48140; *CDEF*, At4g30140; *SYP122*, At3g52400.

### Generation of transgenic lines

The pELTP::mCit-SYP122 construct was obtained by recombining three previously generated entry clones for pELTP(58), mCITRINE and SYP122 cDNA(59) using LR clonase II (Invitrogen). This construct was independently transformed into WT, *sgn3-3*, *myb36-2*, or *sgn3-3 myb36-2* using the floral dip method (60).

### Growth Conditions

For agar plates assays, seeds were surface sterilized, sown on plates containing ½ MS (Murashige and Skoog) with 0.8% agar, stratified for two days at 4°C and grown vertically in growth chamber under long day condition (16h light 100μE 22°C/8h dark 19°C) and observed after 6 days. Piperonylic acid was used from germination at 10μM as described in (7). The CIF2 peptide treatment (DY(SO3H)GHSSPKPKLVRPPFKLIPN) were applied from germination at a concentration of 100 nM. The CIF2 peptide was synthetized by Cambridge Peptided Ltd.

For ionomic analysis, plants were grown using three growth conditions:

- Sterile ½ MS agar plate. Seeds were surface sterilized and sown on plates containing ½ MS (Murashige and Skoog) with 0.8% agar, stratified for two days at 4°C and grown vertically in growth chamber under long day condition (16h light 100μE 22°C/8h dark 19°C). Shoots were collected two weeks after germination.
- Hydroponic. Plants were grown for 5 weeks under short day condition (8h light 100μE 21°C/16h dark 18°C) at 20°C with a relative humidity of 65% RH in a media at pH 5.7 containing 250 μM CaCl_2_, 1 mM KH_2_PO_4_, 50 μM KCl, 250 μM K_2_SO_4_, 1 mM MgSO_4_, 100 μM NaFe-EDTA, 2 mM NH_4_NO_3_, 30 μM H_3_BO_3_, 5 μM MnSO_4_, 1 μM ZnSO_4_, 1 μM CuSO_4_, 0.7 μM NaMoO_4_, 1 μM NiSO_4_. Media was changed weekly.
- Natural soil. Plants were grown for 9 weeks in a growth chamber under short day condition (8h light 100μE 19°C/16h dark 17°C) at 18°C with a relative humidity of 70% in a soil collected in the Sutton Bonington campus of the university of Nottingham (GPS coordinate: 52°49′59.7“N 1°14′56.2“W).

### Fluorescence microscopy

For lignin staining with basic fuchsin, CASP1-GFP visualisation and calclofluor white M2R staining, 6-day-old roots were fixed in paraformaldehyde and cleared in ClearSee as described (61) and using confocal microscopes (Zeiss LSM500 and Leica SP8). Fluorol yellow 088 staining for visualization of suberin was performed and quantified as described in (7, 58) using a fluorescent microscope Leica DM 5000.

### Plasmolysis

Plasmolysis was induced by mounting 6-day-old seedlings in 0.8 M mannitol on microscope slides and directly observed using confocal microscopy (Leica SP8). The proportion of the cell wall length in direct contact with the plasma membrane marker SYP122-mCitrine after plasmolysis was measured using Fiji after plasmolysis. This measurement was done on a maximum projection of the top endodermal cells as seen on Supp. Fig 1B. The quantification represents the percentage of cell wall length in direct contact with the plasma membrane marker SYP122-mCitrine after plasmolysis. Plasmolysis events were imaged and quantified at 15 cells after the onset of elongation.

For the observation of CASP1-GFP and Lignin staining with basic fuchsin, the seedlings were incubated in 0.8 M mannitol for 5 min. and then fixed and cleared as described above.

### Thioacidolysis

The plants were grown for 6 days on ½ MS plates supplemented with 10 nM 6-Benzylaminopurine (BA) and 0.1 % sucrose. Seeds were sown in three parallel lines per square plates (12*12 cm) at high density. Six plates were combined to obtain one replicate. The first 3 mm of root tips as this zone contains no xylem pole were collected in order to obtain 7 to 15 mg of dry weight. The samples were washed twice with 1 mL methanol, rotated for 30 minutes on a carousel and centrifugated to eliminate the methanol supernatant. This washing step was repeated once and the final methanol-extracted samples were then dried for 2 days at 40°C (oven) before thioacidolysis.

The thioacidolyses were carried out in a glass tube with a Teflon-lined screwcap, from about 5 mg sample (weighted at the nearest 0.01 mg) put together with 0.01 mg C21 and 0.01 mg C19 internal standard (50 μL of a 0.2 mg/ml solution) and with 2 ml freshly prepared thioacidolysis reagent. The tightly closed tubes were then heated at 100°C for 4 hours (oil bath), with gentle occasional shaking. After cooling and in each tube, 2 ml of aqueous NaHCO3 0.2M solution were added (to destroy the excess of BF3) and then 0.1 ml HCl 6M (to ensure that the pH is acidic before extraction). The reaction medium was extracted with 2 ml methylene chloride (in the tube) and the lower phase was collected (Pasteur pipette) and dried over Na_2_SO_4_ before evaporation of the solvent (rotoevaporator). The final sample was redissolved in about 2 mL of methylene chloride and 15 μL of this solution were trimethylsilylated (TMS) with 50 μl BSTFA + 5 μl pyridine. The TMS solution was injected (1 μL) into a GC-MS Varian 4000 instrument fitted with an Agilent combiPAL autosampler, a splitless injector (280 °C), and an ion-trap mass spectrometer (electron impact mode, 70 eV), with a source at 220 °C, a transfer line at 280 °C, and an m/z 50−800 scanning range. The GC column was a Supelco SPB1 column (30 m × 0.25 mm i.d., film thickness 0.25 μm) working in the temperature program mode from 45 to 160 °C at +30 °C/min and then 160 to 260 °C at +5 °C/min, with helium as the carrier gas (1 mL/min). The GC-MS determinations of the H, G, and S lignin-derived monomers were carried out on ion chromatograms respectively reconstructed at m/z 239, 269, and 299, as compared to the internal standard hydrocarbon evaluated on the ion chromatogram reconstructed at m/z (57 + 71 + 85).. Each genotype was analyzed as biological triplicates and each biological triplicate was subjected to two different silylations and GC-MS analyses.

### Raman microscopy

Six-day-old seedling were fixed in PBS buffer containing 4% formaldehyde and 1% glutaraldehyde at 4°C overnight, then washed twice with PBS 30 min. Samples were progressively dehydrated in ethanol (30%, 50%, 70%, 100% ethanol). Samples were aligned and embedded in Leica historesin using the protocol described in Beeckman and Viane. 1999. Sections of 5 μm at 4mm from the root tip were generated using a Leica microtome.

The different samples were embedded in resin and cut at 4mm from the root tip using a microtome with a thickness of 15μm. Sections were mounted on superfrost glass slides. The samples were then mapped in a grid over the region of interest. The Raman imaging was performed with a Horiba LabRAM HR spectroscope equipped with a piezoelectric scan stage (Horiba Scientific, UK) using a 532 nm laser, a 100x air objective (Nikon, NA = 0.9) and 600g mm^−1^ grating. Maps were collected for the regions of interest by setting equidistant points along the sample to ensure maximum coverage. The main regions covered in the analysis were the endodermal cell-cell junction, the endodermal cell corners towards the endodermal-cortical junction and the xylem poles (Figure S1). The maps were acquired with 2 accumulations and 30s integration time. The spectra were acquired in the range 300-3100cm^−1^. The spectra were processed using MATLAB and eigenvector software. Firstly, the spectra were trimmed (500-1800 cm^−1^), smoothed and then baseline corrected using an automatic least squares algorithm. This was followed by a percentile mean subtraction (10-20%) to remove signal from the resin. Finally, Gaussian image smoothing was performed to improve signal to noise of the Raman maps. Multivariate Curve Resolution (MCR) analysis was performed on the maps containing the specific ROIs and the corresponding lignin spectra were extracted. These lignin spectra were then used as bounds for the MCR analysis of the large maps where the concentration of these spectra are determined, with a high intensity indicating a high concentration of the specific lignin (Fig. 2C).

### RNA-seq

The plants were grown for 6 days on ½ M/S plates. Seeds were sown in three parallel lines per square plates (12*12 cm) at high density. The first 5 mm of root tips were collected. One plate was used as a biological replicate. The samples were snap-frozen at harvest and ground into fine powder in a 2 mL centrifuge tube. Total RNA was extracted according to Logemann et al., 1987. Samples were homogenized in 400 μL of Z6-buffer containing 8 M guanidine-HCl, 20 mM MES, 20 mM EDTA pH 7.0 After the addition of 400 μl phenol:chloroform:isoamylalcohol, 25:24:1, samples were vortexed and centrifuged (15,000g 10 min.) for phase separation. The aqueous phase was transferred to a new 1.5 mL tube and 0.05 volumes of 1 N acetic acid and 0.7 volumes 96% ethanol was added. The RNA was precipitated at −20°C overnight. Following centrifugation (15,000g 10 min, 4°C), the pellet was washed with 200μL 3M sodium acetate at pH 5.2 and 70% ethanol. The RNA was dried and dissolved in 30 μL of ultrapure water and store at −80°C until use. DNase treatment (DNase I, Amplification Grade, 18068015, Invitrogen) was carried out on the samples to remove genomic DNA. The RNA Concentration and quality were determined using Qubit (Invitrogen; Q10210) and TapeStation (Agilent; G2991A) protocols. Librairies were generated using the Lexogen Quant Seq 3’ mRNA Seq (FWD) Library Prep Kit (Lexogen; 015) which employs polyA selection to enrich for mRNA. Library yield was measured by Qubit (Invitrogen; Q10210) and TapeStation (Agilent; G2991A) systems. protocols to determine concentration and library size, these are then pooled together in equimolar concentrations. The concentration of the pool of libraries were confirmed using the Qubit and qPCR and then loaded onto an Illlumina NextSeq 500/550 High Output Kit v2.5 (75 Cycles) (Illumina; 20024906), to generate approximately 5 million 75bp single-end reads per sample.

Trimmomatic v0.36 was used to identify and discard reads containing the Illumina adaptor sequence. Then, we mapped the resulting high-quality filtered reads against the TAIR10 Arabidopsis reference genome using HISAT2 v.2.1.0 with default parameters. Afterwards, we applied the featureCounts function from the Subread package to count reads that mapped to each one of the 27,206 nuclear protein-coding genes.

We used the R package DESeq2 v.1.24.0 to identify differentially expressed genes (DEGs) between each genotype (*sgn3-3, sgn3-3 myb36-2, sgn3-3 (+CIF2), esb1-1, esb1-1 sgn3-3 and WT (+CIF2*) against WT (Col-0). To do so we fitted the following generalized linear model (GLM).

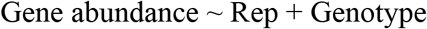

A gene was considered statistically differentially expressed if it had a false discovery rate (FDR) adjusted p-value < 0.1.

For visualization purposes we created a standardized gene matrix. To do so, we applied a variance stabilizing transformation to the raw count gene matrix followed up by standardizing the expression of each gene along the samples. We used this standardized gene matrix to perform principal coordinate (PC) analysis using the prcomp function in R. We displayed the results of the PC analysis using ggplot2.

Additionally, we subset the 3266 statistically significant DEGs from the standardized gene matrix. Then, for each DEG we calculated its mean expression across each genotype followed up by hierarchical clustering (R function hclust method ward.D2) using the euclidean distance for the genotypes and the correlation dissimilarity for the genes. To define the 7 clusters of cohesively expressed genes, we cut the gene dendrogram from the hierarchical clustering using the R function cutree. We visualized the expression of the 3266 DEGs and the result of the clustering approach using ggplot2. We used the compareCluster function from the clusterProfiler R package to perform gene ontology (GO) analysis for the 7 clusters of cohesively expressed DEGs.

We constructed individual heatmaps for the phenylpropanoid pathway and the aquaporin genes by subsetting the corresponding curated gene ids from the standardized gene matrix and procedure described above.

Raw sequence data and read counts are available at the NCBI Gene Expression Omnibus accession number (GEO: GSE158809). Additionally, the scripts created to analyse the RNA-Seq data can be found at https://github.com/isaisg/schengenlignin

### Extraction and profiling of metabolites

The plants (WT, *esb1-1*, *sgn3-3* and *esb1-1 sgn3-3*) were grown for 6 days on ½ M/S plates supplemented with 0.1% sucrose. Seeds were sown in three parallel lines per square plates (12*12 cm) at high density. The first 5 mm of root tips were collected in order to obtain 10 to 20 mg of dry weight per replicate. Eight plates were combined to obtain one replicate. Eight replicates per genotypes were harvested. The samples were snap-frozen at harvest and ground into fine powder in a 2 mL centrifuge tube then homogenized in liquid nitrogen and extracted with 1 ml methanol. The methanol extract was then evaporated, and the pellet dissolved in 200 μl water / cyclohexane (1/1, v/v). 10 μl of the aqueous phase was analyzed via reverse phase UltraHigh Performance Liquid Chromatography (UHPLC; Acquity UPLC Class 1 systems consisting of a Sample Manager-FTN, a Binary Solvent Manager and a Column Manager, Waters Corporation, Milford, MA) coupled to negative ion ElectroSpray Ionization-Quadrupole-Time-of-Flight Mass Spectrometry (ESI-Q-ToF-MS; Vion IMS QTof, Waters Corporation) using an Acquity UPLC BEH C18 column (1.7 μm, 2.1 x 150 mm; Waters Corporation). Using a flow rate of 350 μl/min and a column temperature of 40 °C, a linear gradient was run from 99% aqueous formic acid (0.1%, buffer A) to 50% acetonitrile (0.1% formic acid, buffer B) in 30 min, followed by a further increase to 70% and then to 100% buffer B in 5 and 2 min, respectively. Full MS spectra (m/z 50 – m/z 1,500) were recorded at a scan rate of 10 Hz. The following ESI parameters were used: capillary voltage 2.5 kV, desolvation temperature 550 °C, source temperature 120 °C, desolvation gas 800 L/h and cone gas 50 L/h. Lock correction was applied. In addition to full MS analysis, a pooled sample was subjected to data dependent MS/MS analysis (DDA) using the same separation conditions as above. DDA was performed between m/z 50 and m/z 1,200 at a scan rate of 5 Hz and MS −> MS/MS transition collision energy of 6 eV. The collision energy was ramped from 15 to 35 eV and from 35 to 70 eV for the low and high mass precursor ions, respectively.

Integration and alignment of the m/z features were performed via Progenesis QI software version 2.1 (Waters Corporation). The raw data were imported in this software using a filter strength of 1. A reference chromatogram was manually chosen for the alignment procedure and additional vectors were added in chromatogram regions that were not well aligned. Peak picking was based on all runs with a sensitivity set on ‘automatic’ (value = 5). The normalization was set on ‘external standards’ and was based on the dry weight of the samples (62). In total, 13,091 m/z features were integrated and aligned across all chromatograms. Structural annotation was performed using a retention time window of 1 min, and using both precursor ion and MS/MS identity searches. The precursor ion search (10 ppm tolerance) was based on a compound database constructed via instant JChem (ChemAxon, Budapest, Hungary), whereas MS/MS identities were obtained by matching against an in-house mass spectral database (200 ppm fragment tolerance).

Using R vs 3.4.2., m/z features representing the same compound were grouped following the algorithm in (63). Of the 13,091 m/z features, 12,326 were combined into 2,482 m/z feature groups, whereas 765 remained as m/z feature singlets (i.e. low abundant features). All statistical analyses were performed in R vs. 3.4.2 (64). Including all m/z features and upon applying a prior inverted hyperbolic sine transformation (65), the data were analyzed via both Principal Component Analysis (PCA) and one-way analysis of variance (ANOVA; lm() function) followed by Tukey Honestly Significant Difference (Tukey HSD; TukeyHSD() function) post hoc tests. For PCA, the R packages FactoMineR (66) and factoextra (https://CRAN.R-project.org/package=factoextra) were employed: PCA(scale.unit=T,graph=F), fviz_pca_ind() and fviz_pca_biplot(). Following ANOVA analysis, experiment-wide significant models were revealed via a false discovery rate (FDR) correction using the p.adjust(method=”fdr”) function. Using a FDR-based Q value < 0.05, 4,244 of the 13,091 m/z features were significantly changed in abundance corresponding to 123 m/z feature singlets and 1,158 of the 2,482 compounds. Using a minimum abundance threshold of 500 in at least one of the lines, further analysis was performed on 411 of the 1,158 compounds and 11 of the 123 m/z feature singlets (411 compounds and 11 singlets representing together 889 m/z features).

### Root Hydraulic conductivity

The procedure was exactly identical to the one described in (24). Root hydrostatic conductance (*K*r) was determined in freshly detopped roots using a set of pressure chambers filled with hydroponic culture medium. Excised roots were sealed using dental paste (Coltène/Whaledent s.a.r.l., France) and were subjected to 350 kPa for 10 min to achieve flow stabilization, followed by successive measurements of the flow from the hypocotyl at pressures 320, 160, and 240 kPa. Root hydrostatic conductance (Kr) was calculated by the slope of the flow (*J*v) to pressure relationship. The hydrostatic water conductivity (*L*pr−h, ml H2O g^−1^ h^−1^ MPa^−1^) was calculated by dividing *K*r by the root dry weight.

### Determination of the leaf surface area, mortality and siliques number

For the determination of the leaf surface and mortality, the seeds were stratified for two days at 4°C and the plants were grown in Levington M3 compost in a growth chamber under long day condition (16h light 100μE 21°C/8h dark 19°C). The plants were grown for 7 days with high relative humidity (80%) and then half of the plants were transferred at a lower humidity (65%). Leaf surface was determined at 6, 9, 12 and 15 days after germination using the threshold command of the FiJi software (Schindelin et al., 2012). The plants displaying no growth after 9 days and showing necrosis in all the leave surface were considered as dead plants.

For the determination of the siliques number, the plants were cultivated in a high humidity environment for 10 days after germination and then transferred to a greenhouse. After siliques ripening, only the seeds containing siliques were counted.

### Ionomic analysis with ICP-MS

Ionomics analysis of plants grown in soil (or on plate, hydroponically) was performed as described (67). Briefly, samples (leaf, shoot, root or seed) were harvested into Pyrex test tubes (16 x 100 mm) and dried at 88oC for 20h. After weighing the appropriate number of samples (these masses were used to calculate the rest of the sample masses; alternatively, all samples were weighed individually—usually for small set of samples), the trace metal grade nitric acid Primar Plus (Fisher Chemicals) spiked with indium internal standard was added to the tubes (1 mL per tube). The samples were then digested in dry block heater (DigiPREP MS, SCP Science; QMX Laboratories, Essex, UK) at 115°C for 4 hours. The digested samples were diluted to 10 mL with 18.2 MΩcm Milli-Q Direct water (Merck Millipore). Elemental analysis was performed with an inductively coupled plasma-mass spectrometry (ICP-MS), PerkinElmer NexION 2000 equipped with Elemental Scientific Inc. autosampler, in the collision mode (He). Twenty elements (Li, B, Na, Mg, P, S, K, Ca, Mn, Fe, Co, Ni, Cu, Zn, As, Se, Rb, Sr, Mo and Cd) were monitored. Liquid reference material composed of pooled samples was prepared before the beginning of sample run and was used throughout the whole samples run. It was run after every ninth sample to correct for variation within ICP-MS analysis run (67). The calibration standards (with indium internal standard and blanks) were prepared from single element standards (Inorganic Ventures; Essex Scientific Laboratory Supplies Ltd, Essex, UK) solutions. Sample concentrations were calculated using external calibration method within the instrument software. Further data processing was performed using Microsoft Excel spreadsheet.

## Acknowledgements

We thank Deep Seq (Next Generation Sequencing Facility of the University of Nottingham, UK), the nmRC (Nanoscale and Microscale Research Centre of the University of Nottingham, UK), the Microscopy and Histology Facility of the University of Aberdeen (UK), the VIB Metabolomics Core (VIB-UGent, Belgium). This work was supported by grants from the UK Biotechnology and Biological Sciences Research Council Grant (grant no. BB/N023927/1 to D.E.S.), the Coordinating Action in Plant Sciences Promoting sustainable collaboration in plant sciences (grant no. ERACAPS13.089_RootBarriers to DES), the Engineering and Physical Sciences Research Council (grant no. EP/R025282/1) and the Future Food Beacon of Excellence at the University of Nottingham (Nottingham Research Fellowship to GC, and Postdoctoral Research Fellowship to GR)

## SUPPLEMENTARY FIGURES LEGEND

**Supplemental Figure 1.**
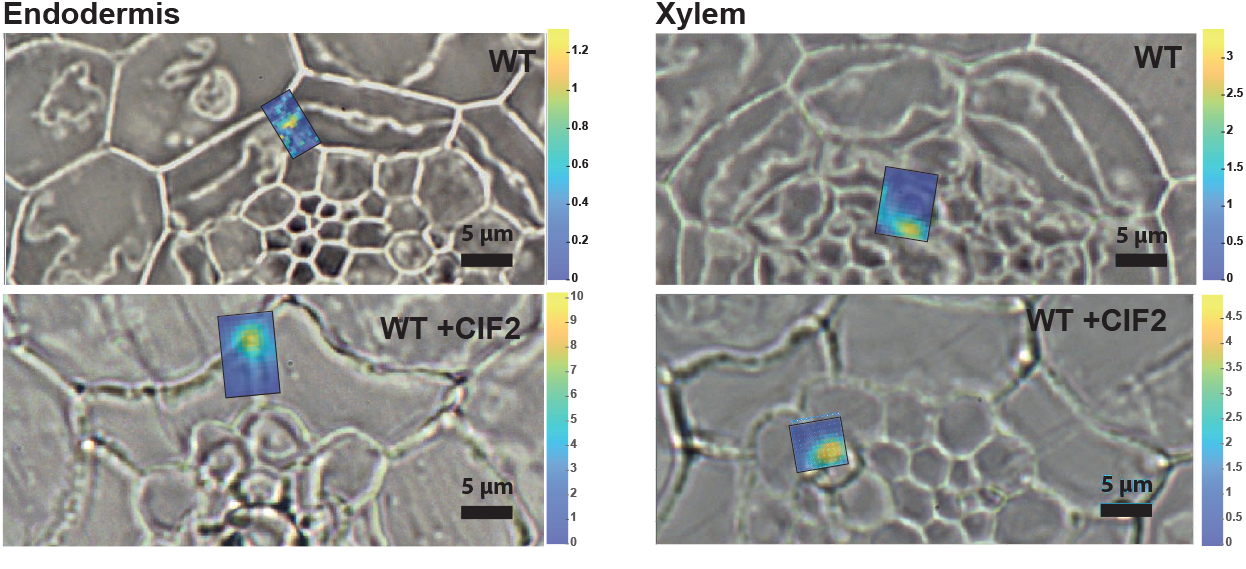
Activation of the Schengen-pathway triggers the deposition of a distinct “stress” lignin in the endodermis. Examples of small Raman maps for endodermal cells of WT(Ø) and WT(+CIF2) and for xylem of WT(Ø) and WT(+CIF2) used for determining the lignin spectra using Multivariate Curve Resolution (MCR) presented in Fig. 2 C-D. The colour code represents the intensity of the lignin factor presented in Fig. 2 C-D.

**Supplemental Figure 2.**
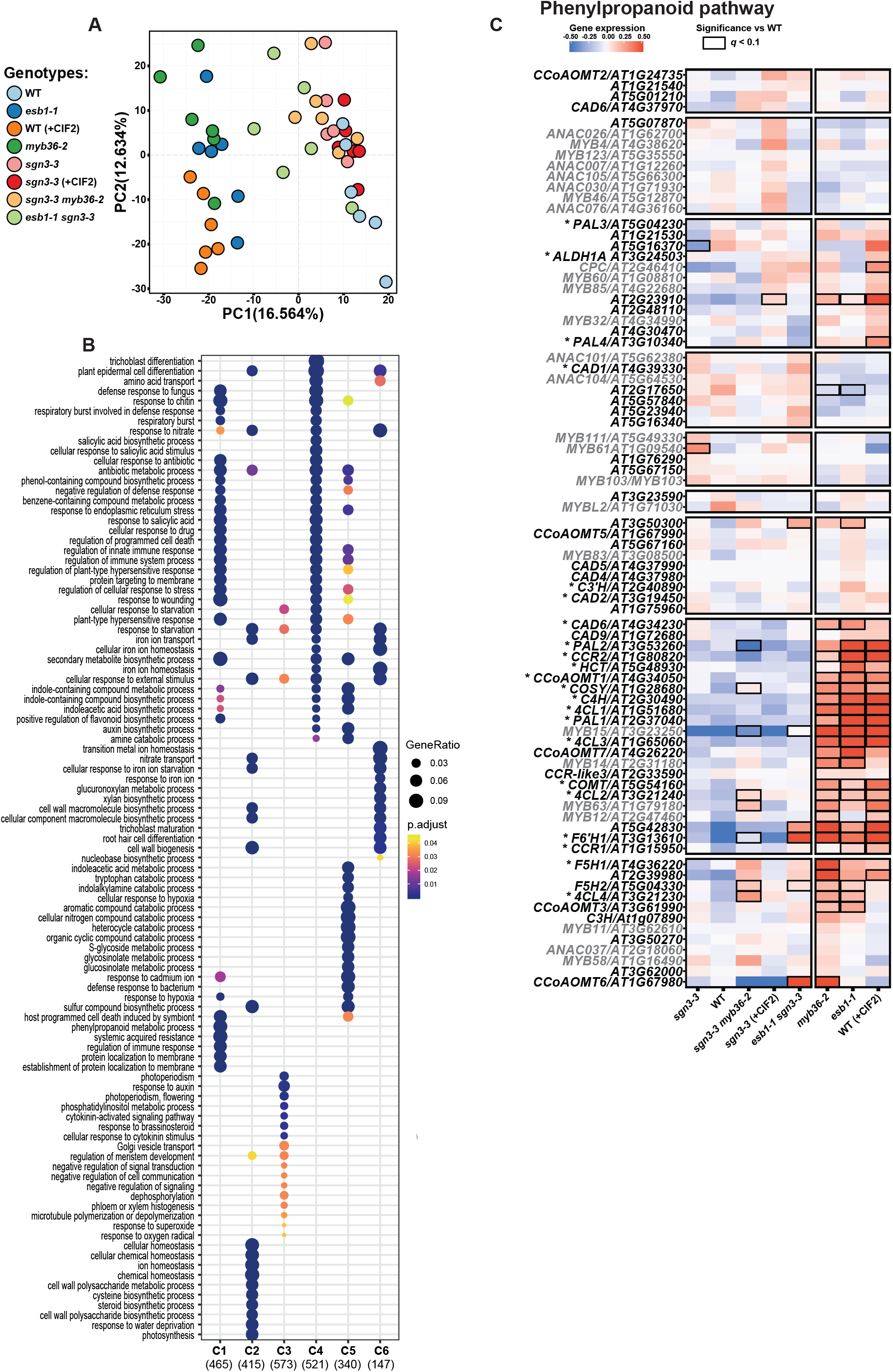
Gene expression profiling in response to the activation of the Schengen-pathway. (**A**) Principal component analysis (PCA) of the differentially expressed genes identified in root tips of wild-type (WT), *sgn3-3*, *esb1-1*, *myb36-2*, *esb1-1 sgn3-3*, *sgn3-3 myb36-2* plants. Treatment with 100 nM CIF2 was applied as indicated (+CIF2) for WT and *sgn3-3* plants (n = 6). (**B**) Gene ontology enrichment in the different gene clusters from Fig. 3A. The colour of each point represents the p-value adjusted using the Benjamin-Hochberg procedure, and the size of each point denotes the percentage of total differential expressed genes in the given gene ontology term (Gene Ratio). (**C**) Heatmap of gene expression of genes related to the phenylpropanoid pathway (black) (69) and their transcriptional regulators (grey) (43, 70). Genes names are given according to (71) for genes related to the phenylpropanoid pathway. Asterisks indicate demonstrated function in lignin biosynthesis with an activity demonstrated *in vitro* or *in vivo* according to (72) for *PAL1-4*, to for (73) *C4H*, (74, 75) for *4CL1-4*, (76, 77) for *CCR1* and *2*, (78, 79) for *CAD1*, 2 and *6*, (80) for *C3’H*, (81) for *C3H*, (82) for *COMT* and *CCoAOMT1*, (83) for *HCT*, (84) for *CSE*, (85) for *ALDH1A*, (45) for *F6’H1*, (46) for *COSY* and (86) for *F5H1*.

**Supplemental Figure 3.**
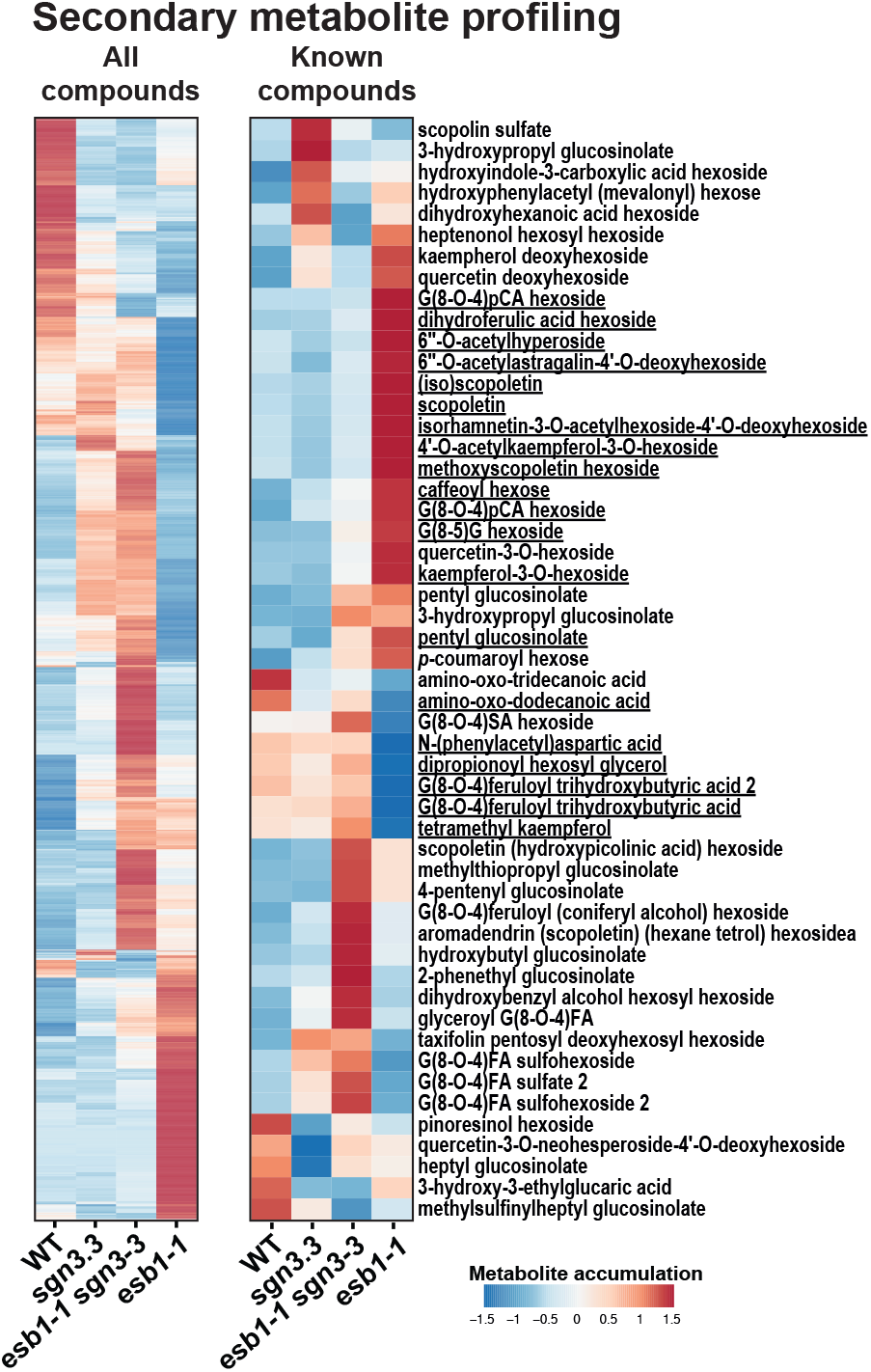
Metabolite profiling in response to the activation of the Schengen-pathway. Heatmaps of metabolite profiling determined using Ultra High Performance Liquid Chromatography (UHPLC) in 5 mm roots tips of wild-type (WT), *sgn3-3*, *esb1-1 sgn3-3* and *esb1-1*. The heatmaps show all the compounds (2497, left) and characterised compounds (52, right) that are differentially accumulated (*q-*value < 0.01, left; *q*-value < 0.1, right n = 8). Underlined names are for compounds that are only differentially accumulated (q-value < 0.1) in *esb1-1* and not changed in *sgn3-3* and *esb1-1 sgn3-3* in comparison with WT. Data for the known compounds are presented in Sup. Table 3.

**Supplemental Figure 4.**
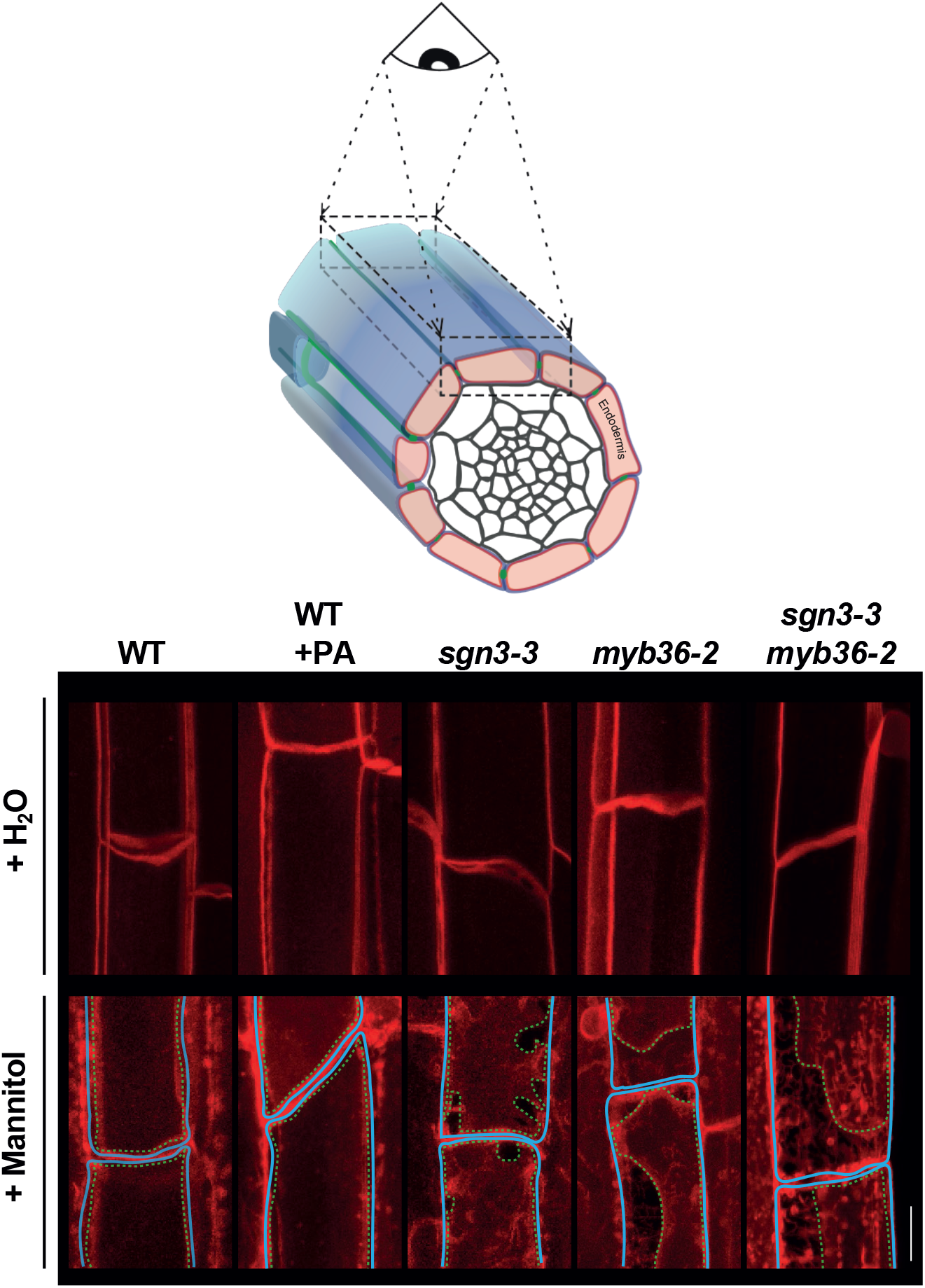
Plasma membrane attachment to the cell wall. (**A**) Maximum projection of the top endodermal cells as shown in the schematic view. The observations were done in lines expressing the plasma membrane marker line pELTP::SYP122mCitrine before plasmolysis (+H2O) and after plasmolysis (+Mannitol) at 15 cells after the onset of elongation. The dashed line represents the contours of the cells. Asterisks show the plasmolysis generated space where no attachment is observed. Scale bar = 5 μm. Representative pictures are shown.

**Supplemental Figure 5.**
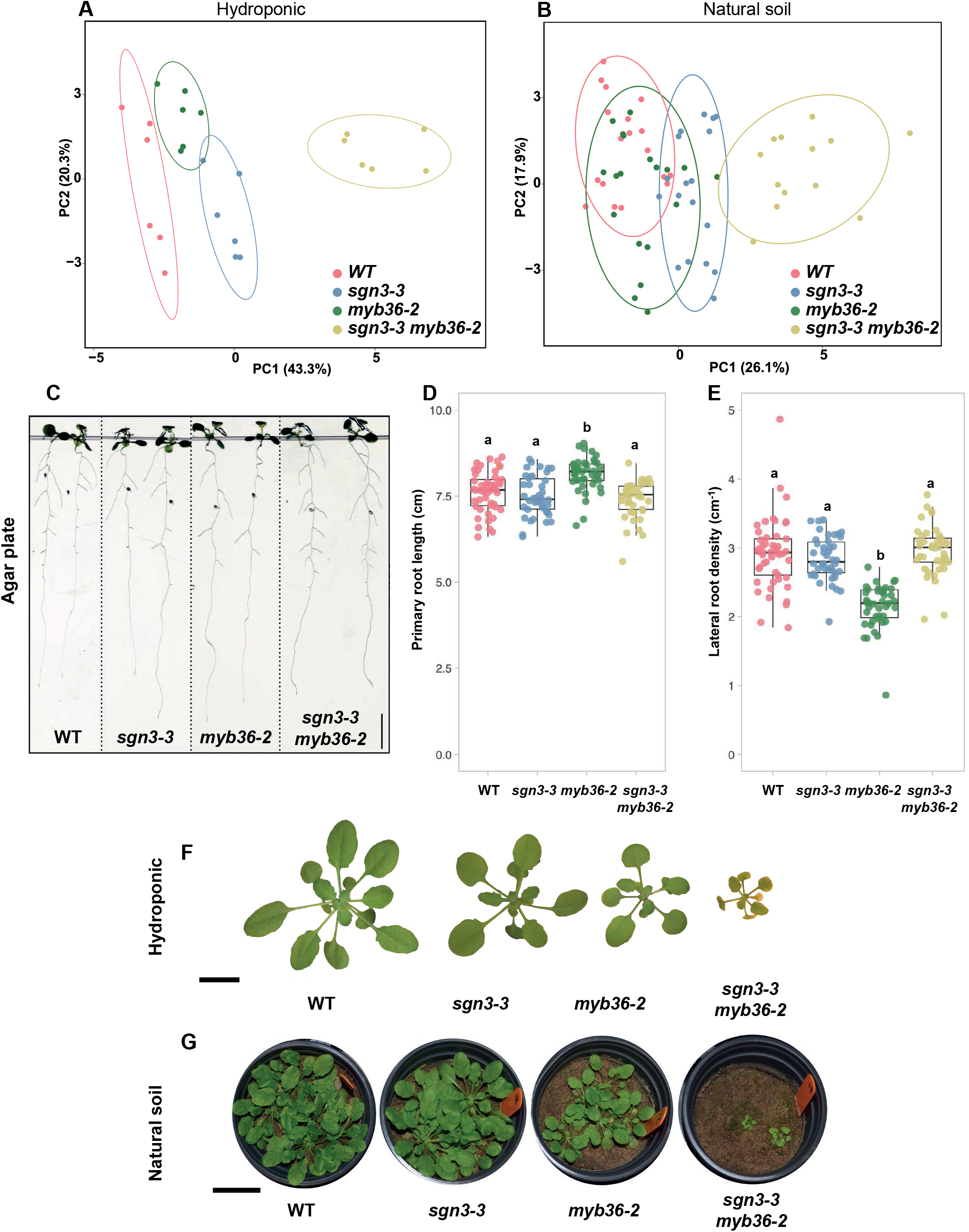
Absence of endodermal apoplastic barrier triggers major ionomic changes in different growth conditions. Principal component analysis (PCA) based on the concentration of 20 elements in shoots of WT, *sgn3-3*, *myb36.3* and *sgn3-3 myb36-2* plants grown in (**A**) hydroponics (short day, n=6) and (**B**) natural soil (short day, n≥13). Ellipses show confidence level at a rate of 90%. (**C**) Pictures of 2-week-old wild-type (WT), *sgn3-3*, *myb36-2* and *sgn3-3 myb36-2* plants grown in agar plates. (**D-E**) Boxplots showing the primary root length (**D**) and lateral roots density (**E**) of 2-week-old WT, *sgn3-3*, *myb36-2* and *sgn3-3 myb36-2* plants grown in agar plates. Letters show significantly different groups according to a Tukey’s test as post hoc analyses (n≥41, P<0.01). (**F**) Pictures of 5-week-old WT, *sgn3-3*, *myb36-2* and *sgn3-3 myb36-2* plants grown in hydroponics. Scale bar = 1 cm. (**G**) Pictures of 9-week-old WT, *sgn3-3*, *myb36-2* and *sgn3-3 myb36-2* plants grown in natural soil. Scale bar = 3 cm.

**Supplemental Figure 6.**
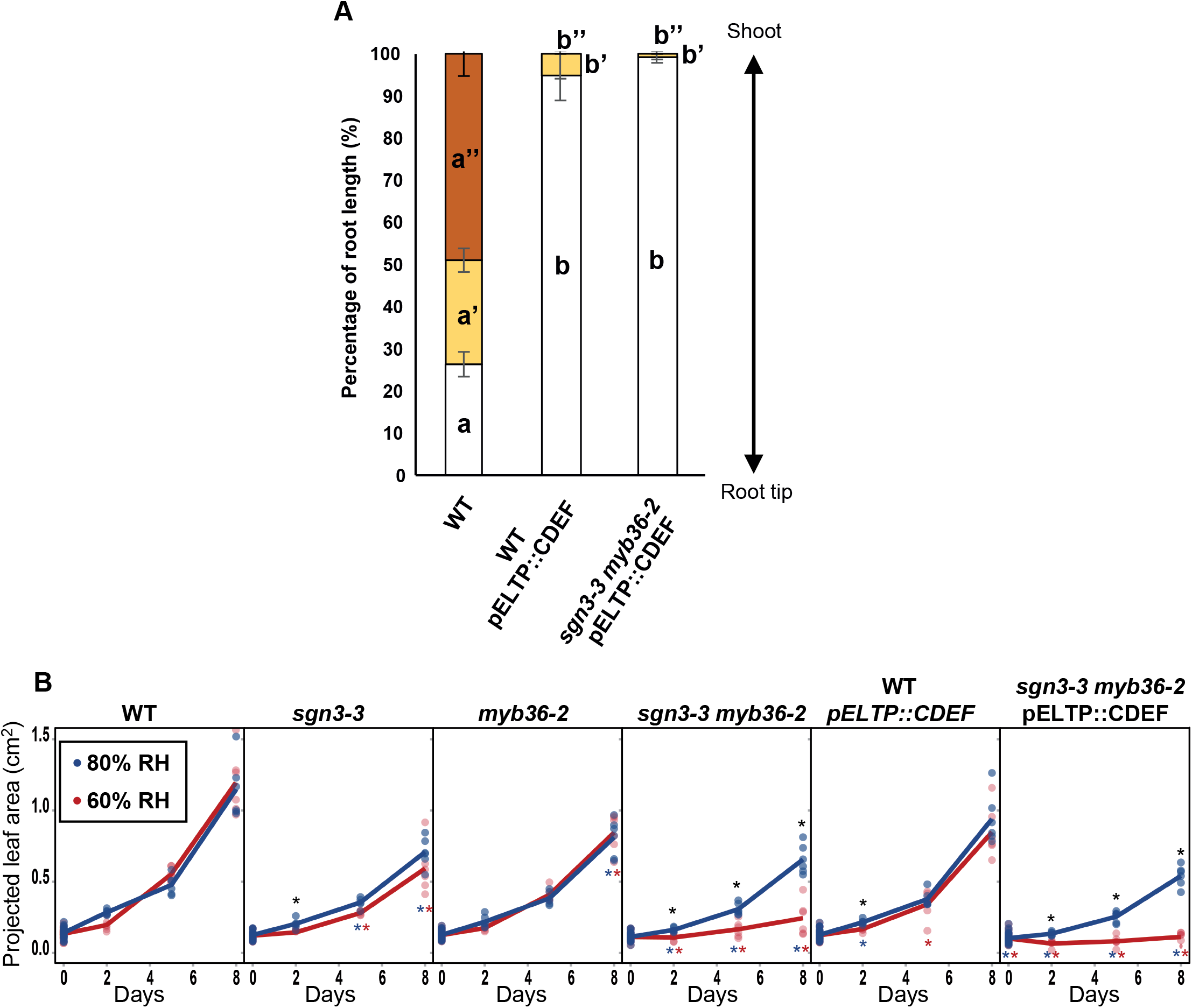
Activation of the Schengen-pathway maintains plant growth under fluctuating environment. (A) Quantification of suberin staining along the root of 6 days-old plants. The results are expressed in percentage of root length divided in three zones: unsuberised (white), discontinuously suberised (yellow), continuously suberised (orange). n = 7, error bars: SD. Individual letters show significant differences using a Mann-Whitney test between the same zones (p<0.01). (**B**) Graphs showing leaf surface area of WT, *sgn3-3*, *myb36-2*, *sgn3-3 myb36-2,* WT-*pELTP::CDEF* and *sgn3-3 myb36-2-pELTP::CDEF* plants germinated in soil with a high humidity (80%) for 7 days and then transferred in an environment with constant (80% RH, blue) or with a lower humidity (60% RH, red). Data were collected 0, 2, 5 and 8 days after the transfer. Each point is the average leave surface per plant from a singles pot (n ≥ 6 pots). Each pot contained at least 6 plants for each genotype. The line shows the average value for each measured time points. Black asterisk indicates a significant difference between high and low humidity for a same genotype at one time point. Blue and red asterisk indicate a significant difference in comparison with WT at the same time point respectively for the high and low humidity environment. The significant differences were calculated using a Tukey’s test as post hoc analyses (p < 0.01).

**Supplemental Table 1. List of the differentially expressed genes in the RNA-seq experiment.**

List of the differentially expressed genes identified in the RNA-seq in root tips of wild-type (WT), *sgn3-3*, *esb1-1*, *myb36-2*, *esb1-1 sgn3-3*, *sgn3-3 myb36-2* plants. Treatment with 100 nM CIF2 was applied as indicated (+CIF2) for WT and *sgn3-3* plants.

**Supplemental Table 2. Metabolite profiling in response to the activation of the Schengen-pathway.**

**Supplemental Table 3. Absence of endodermal apoplastic barrier triggers major ionomic changes.**

Elemental content in shoot of *sgn3-3*, *myb36-2* and *sgn3-3 myb36-2* mutants compared to WT using different growth conditions in agar plates (long day, n=10), in hydroponics (short day, n=6) and natural soil (short day, n≥13). Elements concentration were determined by ICP-MS. Data are presented as mean ± standard deviation (SD). *t tests* were performed to determine the significant differences to WT and the corresponding p-values are presented.

## References

1. Boerjan W, Ralph J, Baucher M (2003) Lignin biosynthesis. Annu Rev Plant Biol 54:519–546.

2. Lee MH, et al. (2019) Lignin-based barrier restricts pathogens to the infection site and confers resistance in plants. The EMBO Journal 38(23):e1745–17.

3. Vanholme R, De Meester B, Ralph J, Boerjan W (2019) Lignin biosynthesis and its integration into metabolism. Current Opinion in Biotechnology 56:230–239.

4. Tobimatsu Y, Schuetz M (2018) Lignin polymerization: how do plants manage the chemistry so well? Current Opinion in Biotechnology 56:75–81.

5. Schuetz M, et al. (2014) Laccases direct lignification in the discrete secondary cell wall domains of protoxylem. PLANT PHYSIOLOGY 166(2):798–807.

6. Barros J, Serk H, Granlund I, Pesquet E (2015) The cell biology of lignification in higher plants. Annals of Botany:mcv046.

7. Naseer S, et al. (2012) Casparian strip diffusion barrier in Arabidopsis is made of a lignin polymer without suberin. Proc Natl Acad Sci USA 109(25):10101–10106.

8. Geldner N (2013) The endodermis. Annu Rev Plant Biol 64(1):531–558.

9. Pfister A, et al. (2014) A receptor-like kinase mutant with absent endodermal diffusion barrier displays selective nutrient homeostasis defects. Elife 3. doi:10.7554/eLife.03115.

10. Baxter I, et al. (2012) Biodiversity of Mineral Nutrient and Trace Element Accumulation in Arabidopsis thaliana. PLoS ONE 7(4):e35121.

11. Barbosa ICR, Rojas-Murcia N, Geldner N (2018) The Casparian strip—one ring to bring cell biology to lignification? Current Opinion in Biotechnology 56:121–129.

12. Roppolo D, et al. (2011) A novel protein family mediates Casparian strip formation in the endodermis. Nature 473(7347):380–383.

13. Rojas-Murcia N, et al. (2020) High-order mutants reveal an essential requirement for peroxidases but not laccases in Casparian strip lignification. bioRxiv:2020.06.17.154617.

14. Lee Y, Rubio MC, Alassimone J, Geldner N (2013) A Mechanism for Localized Lignin Deposition in the Endodermis. Cell 153(2):402–412.

15. Hosmani PS, et al. (2013) Dirigent domain-containing protein is part of the machinery required for formation of the lignin-based Casparian strip in the root. Proc Natl Acad Sci USA 110(35):14498–14503.

16. Liberman LM, Sparks EE, Moreno-Risueno MA, Petricka JJ, Benfey PN (2015) MYB36 regulates the transition from proliferation to differentiation in the Arabidopsis root. Proc Natl Acad Sci USA:201515576.

17. Kamiya T, et al. (2015) The MYB36 transcription factor orchestrates Casparian strip formation. Proc Natl Acad Sci USA:201507691.

18. Fujita S, et al. (2020) SCHENGEN receptor module drives localized ROS production and lignification in plant roots. The EMBO Journal:e103894.

19. Nakayama T, et al. (2017) A peptide hormone required for Casparian strip diffusion barrier formation in Arabidopsis roots. Science 355(6322):284–286.

20. Doblas VG, et al. (2017) Root diffusion barrier control by a vasculature-derived peptide binding to the SGN3 receptor. Science 355(6322):280–284.

21. Alassimone J, et al. (2016) Polarly localized kinase SGN1 is required for Casparian strip integrity and positioning. Nature Plants:1–10.

22. Li B, et al. (2017) Role of LOTR1 in Nutrient Transport through Organization of Spatial Distribution of Root Endodermal Barriers. Curr Biol:1–9.

23. Alassimone J, Naseer S, Geldner N (2010) A developmental framework for endodermal differentiation and polarity. Proc Natl Acad Sci USA 107(11):5214–5219.

24. Wang P, et al. (2019) Surveillance of cell wall diffusion barrier integrity modulates water and solute transport in plants. Scientific Reports 9(1):4227.

25. Agarwal UP, McSweeny JD, Ralph SA (2011) FT–Raman Investigation of Milled-Wood Lignins: Softwood, Hardwood, and Chemically Modified Black Spruce Lignins. Journal of Wood Chemistry and Technology 31(4):324–344.

26. Mahonen AP (2006) Cytokinin Signaling and Its Inhibitor AHP6 Regulate Cell Fate During Vascular Development. Science 311(5757):94–98.

27. Lange BM, Lapierre C, Sandermann H Jr (1995) Elicitor-Induced Spruce Stress Lignin (Structural Similarity to Early Developmental Lignins). Plant Physiol 108(3):1277.

28. Fukushima K, Terashima N (1991) Heterogeneity in formation of lignin. Wood Science and Technology 25(5):371–381.

29. Westermark U (1985) The occurrence of p-hydroxyphenylpropane units in the middle-lamella lignin of spruce (Picea abies). Wood Science and Technology 19(3):223–232.

30. Lapierre C (1995) Application of New Methods for the Investigation of Lignin Structure. Forage Cell Wall Structure and Digestibility (John Wiley & Sons, Ltd), pp 133–166.

31. Ride JP (1975) Lignification in wounded wheat leaves in response to fungi and its possible rôle in resistance. Physiological Plant Pathology 5(2):125–134.

32. Hammerschmidt R, Bonnen AM, Bergstrom GC, Baker KK (1985) Association of epidermal lignification with nonhost resistance of cucurbits to fungi. Can J Bot 63(12):2393–2398.

33. Doster MA, Bostock RM (1988) Quantification of lignin formation in almond bark in response to wounding and infection by Phytophthora species. Phytopathology 78(4):473–477.

34. Lange BM, Lapierre C, (null) HSJP (1995) Elicitor-induced spruce stress lignin (structural similarity to early developmental lignins). PLANT PHYSIOLOGY. doi:10.1007/BF00029715.

35. Campbell MM, Ellis BE (1992) Fungal elicitor-mediated responses in pine cell cultures: cell wall-bound phenolics*. The International Journal of Plant Biochemistry 31(3):737–742.

36. Bonawitz ND, et al. (2014) Disruption of Mediator rescues the stunted growth of a lignin-deficient Arabidopsis mutant. Nature 509(7500):376–380.

37. Franke R, et al. (2002) Changes in secondary metabolism and deposition of an unusual lignin in the ref8 mutant of Arabidopsis. Plant J 30(1):47–59.

38. Coleman HD, Park J-Y, Nair R, Chapple C, Mansfield SD (2008) RNAi-mediated suppression of p-coumaroyl-CoA 3′-hydroxylase in hybrid poplar impacts lignin deposition and soluble secondary metabolism. Proc Natl Acad Sci USA 105(11):4501–4506.

39. Ralph J, Akiyama T, Coleman HD, Mansfield SD (2012) Effects on Lignin Structure of Coumarate 3-Hydroxylase Downregulation in Poplar. Bioenerg Res 5(4):1009–1019.

40. Rogers LA, et al. (2005) Comparison of lignin deposition in three ectopic lignification mutants. New Phytol 168(1):123–140.

41. Hématy K, et al. (2007) A Receptor-like Kinase Mediates the Response of Arabidopsis Cells to the Inhibition of Cellulose Synthesis. Current Biology 17(11):922–931.

42. Cheung AY, Wu H-M (2011) THESEUS 1, FERONIA and relatives: a family of cell wall-sensing receptor kinases? Current Opinion in Plant Biology 14(6):632–641.

43. Liu J, Osbourn A, Ma P (2015) MYB Transcription Factors as Regulators of Phenylpropanoid Metabolism in Plants. Molecular Plant 8(5):689–708.

44. Chezem WR, Memon A, Li F-S, Weng J-K, Clay NK (2017) SG2-Type R2R3-MYB Transcription Factor MYB15 Controls Defense-Induced Lignification and Basal Immunity in Arabidopsis. THE PLANT CELL ONLINE 29(8):1907–1926.

45. Kai K, et al. (2008) Scopoletin is biosynthesized via ortho-hydroxylation of feruloyl CoA by a 2-oxoglutarate-dependent dioxygenase in Arabidopsis thaliana. The Plant Journal 55(6):989–999.

46. Vanholme R, et al. (2019) COSY catalyses trans–cis isomerization and lactonization in the biosynthesis of coumarins. Nature Plants:1–12.

47. Stringlis IA, et al. (2018) MYB72-dependent coumarin exudation shapes root microbiome assembly to promote plant health. Proc Natl Acad Sci USA 115(22):E5213.

48. Zhalnina K, et al. (2018) Dynamic root exudate chemistry and microbial substrate preferences drive patterns in rhizosphere microbial community assembly. Nature Microbiology 3(4):470–480.

49. König S, et al. (2014) Soluble phenylpropanoids are involved in the defense response of Arabidopsis against Verticillium longisporum. New Phytol 202(3):823–837.

50. Kroemer K (1903) Wurzelhaut, Hypodermis und Endodermis der Angiospermenwurzel (Bibl Bot).

51. Behrisch R (1926) Zur Kenntnis der Endodermiszelle. Berichte der Deutschen Botanischen Gesellschaft 44(*3*):162–164.

52. Fernández-Marcos M, et al. (2016) Control of Arabidopsis lateral root primordium boundaries by MYB36. New Phytol:1–8.

53. Javot H, Maurel C (2002) The role of aquaporins in root water uptake. Annals of Botany 90(3):301–313.

54. Javot H, et al. (2003) Role of a single aquaporin isoform in root water uptake. Plant Cell 15(2):509–522.

55. Alexandersson E, et al. (2005) Whole gene family expression and drought stress regulation of aquaporins. Plant Mol Biol 59(3):469–484.

56. Olas JJ, Fichtner F, Apelt F (2019) All roads lead to growth: imaging-based and biochemical methods to measure plant growth. Journal of Experimental Botany 71(1):11–21.

57. Andersen TG, et al. (2018) Diffusible repression of cytokinin signalling produces endodermal symmetry and passage cells. Nature:1–20.

58. Barberon M, et al. (2016) Adaptation of Root Function by Nutrient-Induced Plasticity of Endodermal Differentiation. Cell:1–35.

59. Vermeer JEM, et al. (2014) A Spatial Accommodation by Neighboring Cells Is Required for Organ Initiation in Arabidopsis. Science 343(6167):178–183.

60. Clough SJ, Bent AF (1998) Floral dip: a simplified method for Agrobacterium-mediated transformation of Arabidopsis thaliana. Plant J 16(6):735–743.

61. Ursache R, Andersen TG, Marhavy P, Geldner N (2018) A protocol for combining fluorescent proteins with histological stains for diverse cell wall components. The Plant Journal 93(2):399–412.

62. Morreel K, et al. (2006) Genetical metabolomics of flavonoid biosynthesis in Populus: a case study. The Plant Journal 47(2):224–237.

63. Morreel K, et al. (2014) Systematic structural characterization of metabolites in Arabidopsis via candidate substrate-product pair networks. THE PLANT CELL ONLINE 26(3):929–945.

64. R: A language and environment for statistical computing. R Foundation for Statistical Computing, Vienna, Austria. (2018) R: A language and environment for statistical computing. R Foundation for Statistical Computing, Vienna, Austria. R Core Team. Available at: https://www.R-project.org/.

65. Burbidge JB, Magee L, Robb AL (1988) Alternative Transformations to Handle Extreme Values of the Dependent Variable. Journal of the American Statistical Association 83(401):123–127.

66. Lê S, Josse J, Husson F (2008) FactoMineR: An R Package for Multivariate Analysis. Journal of Statistical Software; Vol 1, Issue 1 (2008).

67. Danku JMC, Lahner B, Yakubova E, Salt DE (2013) Large-scale plant ionomics. Methods Mol Biol 953(Chapter 17):255–276.

68. Vanholme B, Houari El I, Boerjan W (2018) Bioactivity: phenylpropanoids’ best kept secret. Current Opinion in Biotechnology 56:156–162.

69. Mueller LA, Zhang P, Rhee SY (2003) AraCyc: a biochemical pathway database for Arabidopsis. PLANT PHYSIOLOGY 132(2):453–460.

70. Ohtani M, Demura T (2018) The quest for transcriptional hubs of lignin biosynthesis: beyond the NAC-MYB-gene regulatory network model. Current Opinion in Biotechnology 56:82–87.

71. Raes J, Rohde A, Christensen JH, Van de Peer Y, Boerjan W (2003) Genome-wide characterization of the lignification toolbox in Arabidopsis. PLANT PHYSIOLOGY 133(3):1051–1071.

72. Huang J, et al. (2010) Functional Analysis of the Arabidopsis *PAL* Gene Family in Plant Growth, Development, and Response to Environmental Stress. Plant Physiol 153(4):1526.

73. Schilmiller AL, et al. (2009) Mutations in the cinnamate 4-hydroxylase gene impact metabolism, growth and development in Arabidopsis. The Plant Journal 60(5):771–782.

74. Costa MA, et al. (2005) Characterization in vitro and in vivo of the putative multigene 4-coumarate:CoA ligase network in Arabidopsis: syringyl lignin and sinapate/sinapyl alcohol derivative formation. The International Journal of Plant Biochemistry 66(17):2072–2091.

75. Li Y, Kim JI, Pysh L, Chapple C (2015) Four Isoforms of Arabidopsis 4-Coumarate:CoA Ligase Have Overlapping yet Distinct Roles in Phenylpropanoid Metabolism. PLANT PHYSIOLOGY 169(4):2409–2421.

76. Lauvergeat V, et al. (2001) Two cinnamoyl-CoA reductase (CCR) genes from Arabidopsis thaliana are differentially expressed during development and in response to infection with pathogenic bacteria. The International Journal of Plant Biochemistry 57(7):1187–1195.

77. Baltas M, et al. (2005) Kinetic and inhibition studies of cinnamoyl-CoA reductase 1 from Arabidopsis thaliana. Plant Physiology et Biochemistry 43(8):746–753.

78. Sibout R, et al. (2005) CINNAMYL ALCOHOL DEHYDROGENASE-C and -D are the primary genes involved in lignin biosynthesis in the floral stem of Arabidopsis. THE PLANT CELL ONLINE 17(7):2059–2076.

79. Eudes A, et al. (2006) Evidence for a role of AtCAD 1 in lignification of elongating stems of Arabidopsis thaliana. Planta 225(1):23–39.

80. Franke R, et al. (2002) The Arabidopsis REF8 gene encodes the 3-hydroxylase of phenylpropanoid metabolism. Plant J 30(1):33–45.

81. Barros J, et al. (2019) 4-Coumarate 3-hydroxylase in the lignin biosynthesis pathway is a cytosolic ascorbate peroxidase. Nature Communications:1–11.

82. Do C-T, et al. (2007) Both caffeoyl Coenzyme A 3-O-methyltransferase 1 and caffeic acid O-methyltransferase 1 are involved in redundant functions for lignin, flavonoids and sinapoyl malate biosynthesis in Arabidopsis. Planta 226(5):1117–1129.

83. Hoffmann L, Maury S, Martz F, Geoffroy P, Legrand M (2003) Purification, cloning, and properties of an acyltransferase controlling shikimate and quinate ester intermediates in phenylpropanoid metabolism. J Biol Chem 278(1):95–103.

84. Vanholme R, et al. (2013) Caffeoyl shikimate esterase (CSE) is an enzyme in the lignin biosynthetic pathway in Arabidopsis. Science 341(6150):1103–1106.

85. Nair RB, Bastress KL, Ruegger MO, Denault JW, Chapple C (2004) The Arabidopsis thaliana REDUCED EPIDERMAL FLUORESCENCE1 gene encodes an aldehyde dehydrogenase involved in ferulic acid and sinapic acid biosynthesis. THE PLANT CELL ONLINE 16(2):544–554.

86. Meyer K, Shirley AM, Cusumano JC, Bell-Lelong DA, Chapple C (1998) Lignin monomer composition is determined by the expression of a cytochrome P450-dependent monooxygenase in Arabidopsis. Proceedings of the National Academy of Sciences 95(12):6619–6623.

